# Multiple pathways couple kinetochore orientation to the meiotic spindle cycle

**DOI:** 10.64898/2026.04.17.719180

**Authors:** Aparna Vinod, Matthew Turner, Shaun Webb, Adele L. Marston

## Abstract

Meiosis generates haploid gametes from a diploid progenitor. In meiosis I, homologous chromosomes segregate while sister chromatids co-orient toward the same spindle pole. The mechanisms underlying this specialized segregation pattern remain incompletely understood. Here we identify pathways required for meiosis I chromosome segregation through a forward genetic screen in *Saccharomyces cerevisiae*. We find that meiosis I is highly sensitive to perturbations in diverse components of the segregation machinery and define key functional interfaces within complexes important for this division. These include meiosis I–specific regulators that establish the specialized chromosome pattern, namely the monopolin complex that directs sister kinetochore co-orientation and the Spo13^MOKIR^–Cdc5^Polo^ module that promotes meiosis I chromosome segregation. In addition, we identify mutations affecting the core segregation machinery, including the spindle pole body, spindle midzone, and outer kinetochore, which can disrupt coupling between the chromosome segregation program and the meiotic spindle cycle. Together, our findings reveal that multiple pathways coordinate chromosome segregation with the meiotic divisions and highlight the unique demands of meiosis I.

## Introduction

Meiosis generates haploid gametes through two successive chromosome segregation events following a single round of DNA replication. Maternal and paternal chromosomes, known as homologs, are segregated in meiosis I while sister chromatids are segregated in meiosis II (Duro and Marston, 2015). Accurate segregation of homologs in meiosis I requires physical linkage between homologous chromosomes, typically provided by chiasmata formed during meiotic crossover recombination, together with cohesion along chromosome arms. Since sister kinetochores are mono-oriented in meiosis I and attach to microtubules from the same pole, these linkages allow homolog pairs to generate tension on the meiosis I spindle. At anaphase I onset, cohesion is lost from chromosome arms, triggering homolog segregation. However, cohesion is retained in pericentromeric regions until meiosis II when sister kinetochores are bioriented and attach to microtubules from opposite poles to generate tension between sister chromatids.

How sister kinetochore mono-orientation is established in meiosis I remains incompletely understood. In budding yeast *Saccharomyces cerevisiae*, the monopolin complex comprising the meiosis-specific Mam1 protein, together with Csm1, Lrs4 and the casein kinase Hrr25 (CK1δ) is essential for monoorientation (Toth et al., 2000; Rabitsch et al., 2003; Petronczki et al., 2006). Monopolin forms a V-shaped structure that binds an N-terminal extension on the Dsn1 kinetochore subunit and is thought to fuse the two sister kinetochores together into a single microtubule-binding unit (Sarangapani et al., 2014; Sarkar et al., 2013; Plowman et al., 2019; Corbett and Harrison, 2016; Corbett et al., 2010). However, there is no evidence that monopolin-like proteins are important for mono-orientation outside budding yeast. Instead, cohesion and a family of proteins collectively known as <u>M</u>eiosis <u>O</u>ne <u>Ki</u>nase <u>R</u>egulators (MOKIRs) appear to be broadly required for mono-orientation, including in *S. cerevisiae* (Chelysheva et al., 2005; Parra et al., 2004; Sakuno et al., 2009; Severson et al., 2009; Barton et al., 2022; Singh et al., 2025; Kim et al., 2015; Yokobayashi and Watanabe, 2005). Like other MOKIR proteins, *S. cerevisiae* Spo13 associates with the Polo-like kinase Cdc5 via its Polo-binding domain (PBD), which is itself required for sister kinetochore mono-orientation (Matos et al., 2008; Galander et al., 2019; Lee and Amon, 2003; Clyne et al., 2003). Cdc5^Plk1^ works to promote mono-orientation together with the Dbf4-dependent kinase, Cdc7 (DDK), at least in part through regulation of monopolin subunit Lrs4 (Matos et al., 2008; Chen and Weinreich, 2010). MOKIRs also contribute to other aspects of the meiotic segregation pattern, including the regulation of pericentromeric cohesion protection and the execution of two successive divisions (Kim et al., 2015; Yokobayashi and Watanabe, 2005; Lee et al., 2004; Katis et al., 2004).

In addition to establishing the correct chromosome configuration, meiotic segregation must occur in coordination with the meiotic spindle cycle. Following meiosis I, cells must disassemble the first spindle, duplicate spindle pole bodies and assemble a second spindle to drive meiosis II. In budding yeast, these events are regulated by the FEAR (Cdc14 Early Anaphase Release) pathway, which promotes transient activation of the Cdc14 phosphatase during anaphase I (Marston et al., 2003; Buonomo et al., 2003). In the absence of FEAR activity, cells fail to duplicate spindle pole bodies following meiosis I (Fox et al., 2017; Bizzari and Marston, 2011). Nevertheless, cells attempt to assemble a second spindle along the same axis at the time meiosis II would normally occur (Bizzari and Marston, 2011). Because other cell-cycle events continue, including the switch to bioriented kinetochores, a fraction of sister chromatids segregate on this reassembled spindle (Bizzari and Marston, 2011). As a consequence of this mixed segregation, aneuploid dyads are produced instead of tetrads. This phenotype is striking because it reveals that progression through the meiotic division program can proceed despite disruption of the spindle cycle, suggesting that successful meiosis I requires particularly robust coordination between chromosome segregation and the meiotic cell cycle.

This uncoupling of chromosome segregation from the meiotic spindle cycle provides a sensitised system to identify pathways that ensure chromosomes segregate during the correct meiotic division. Indeed, *spo12Δ* mutants have previously been used as the basis of a genetic screen that identified components of the monopolin complex required for sister kinetochore mono-orientation (Rabitsch et al., 2003). Similar screens in *Schizosaccharomyces pombe* and *Arabidopsis thaliana* have also identified genes important for mono-orientation (Singh et al., 2025; Yokobayashi and Watanabe, 2005). A limitation of the *S. cerevisiae* screen is that it relied on transposon mutagenesis (Rabitsch et al., 2003), which primarily disrupts non-essential genes and therefore cannot reveal functions of essential proteins or define functional protein interfaces. To overcome these limitations, we adapted this strategy to perform a forward genetic screen using EMS mutagenesis in budding yeast. Our goal was to identify mutations that restore spore viability in *spo12Δ* cells and thereby uncover pathways that coordinate kinetochore orientation and chromosome segregation with the meiotic spindle cycle during meiosis I.

## Results

### A genetic screen identifies point mutations affecting meiosis I

To identify mutations that cause sister chromatids to biorient rather than mono-orient in meiosis, we adapted a previously described strategy (Rabitsch et al., 2003). The screen exploits the fact that spore inviability after single-division meiosis arises from random homolog segregation and can be rescued by forcing even segregation of sister chromatids. Single-division meiosis is achieved using a *spo12Δ* background in which the second round of spindle pole body duplication is prevented, resulting in the formation of dyads (two spores) rather than tetrads (four spores) (Marston et al., 2003; Buonomo et al., 2003; Fox et al., 2017; Klapholz and Esposito, 1980). To prevent homolog linkage via chiasmata, we used *spo11Δ* which blocks the initiation of meiotic recombination causing random segregation of homologs in meiosis I (Keeney et al., 1997; Shonn et al., 2000). We reasoned that even chromosome segregation, and consequently spore viability, could be rescued in two principal ways in *spo11Δ spo12Δ* cells. First, disruption of mono-orientation, for example in *mam1Δ*, would force sister chromatids to bi-orient and segregate evenly in meiosis I. Alternatively, mutations that prevent chromosomes from being pulled apart by the spindle in meiosis I could delay segregation until kinetochores have already undergone a meiosis II-like switch to biorientation. The first class of mutations would therefore identify factors that direct mono-orientation, whereas the second class of mutations would reveal general chromosome segregation pathways that are particularly critical for the specialised meiosis I division (Figure 1A).

**Figure 1.**
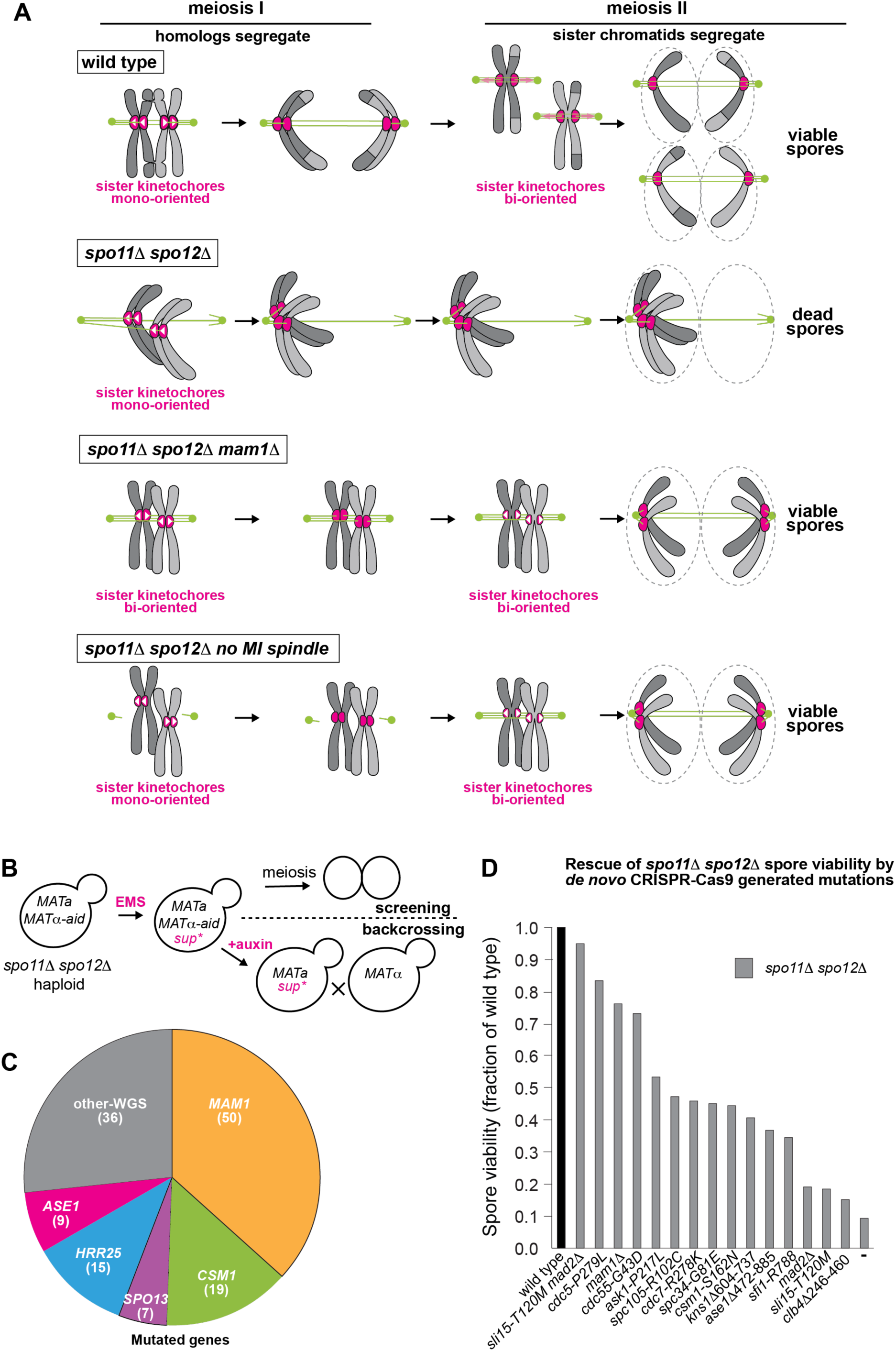
A genetic screen to identify genes that coordinate kinetochore orientation with chromosome segregation in *S. cerevisiae*. (A) Schematic explaining the rationale behind the screen. In wild type, homologs segregate in meiosis I due to sister kinetochore mono-orientation and chiasmata. Spindle pole bodies (SPBs) then re-duplicate to support a second round of chromosome segregation in meiosis II, where sister kinetochores are now bi-oriented. In the single division of *spo11Δ spo12Δ* cells, sister kinetochores mono-orient but homologs segregate randomly to produce inviable spores. In *spo11Δ spo12Δ mam1Δ* cells, sister kinetochores bi-orient already in meiosis I, leading to even segregation and viable spores. If meiosis I chromosome segregation is prevented in the *spo11Δ spo12Δ* background, *e.g.* by disrupting spindle formation, mono-orientation is lost prior to kinetochore capture by microtubules resulting in even sister chromatid segregation at the time of meiosis II and the formation of viable spores. Therefore, isolation of mutations that rescue the spore inviability of *spo11Δ spo12Δ* cells will identify key pathways important for mono-orientation and its co-ordination with the cell cycle. (B) Screening strategy through design of a conditional pseudodiploid to allow screening and backcrossing for mutation identification. Haploid *spo11Δ spo12Δ* cells were engineered to express both *MATa* and *MATα* mating information, which is essential for sporulation/meiosis. Both MATα1 and MATα2 were fused to the auxin inducible degron (aid) so that following EMS mutagenesis and selection of viable spores, MATα information can be degraded to allow mating to unmutagenised *MATa* cells, reducing the penetrance of non-causal mutations in progeny by half. (C) Overview of genes identified in the screen. Numbers represent the number of different isolates with mutations in the indicated genes, identified either through complementation analyses or whole genome sequencing. (D) Confirmation of mutation causality. Mutations were generated *de novo* and introduced into *spo11Δ spo12Δ* pseudodiploid cells. Spore viability following random spore analysis is shown as a fraction of wild type for the indicated strains.

This strategy was previously used with transposon-directed mutagenesis to identify components of monopolin (Rabitsch et al., 2003), but transposon disruption primarily identifies non-essential genes and therefore misses essential factors required for meiosis I. To overcome this limitation and enable identification of functional protein surfaces, we adapted the approach to generate point mutations using the chemical mutagen Ethyl methanesulfonate (EMS). Because EMS introduces multiple mutations throughout the genome, recovery of causal mutations requires backcrossing to remove unrelated background mutations. To enable this while performing the screen in haploids, we engineered a conditional system in which *MATa* cells express *MATα* regulators to permit meiosis but regain mating competence upon auxin (NAA) addition (Figure 1B). Both M*ATα* regulators, *MATα1* and *MATα2*, were fused to an auxin-inducible degron in haploid cells expressing OsTIR1, enabling auxin-dependent degradation and restoration of mating (Figure S1A-C; see Materials and Methods).

Following two rounds of EMS mutagenesis, we recovered 136 isolates with improved spore viability compared to the *spo11Δ spo12Δ* control, and were able to assign likely causative genes to 123 of them (Figure 1C; Table S1). We used complementation testing to determine whether causal mutations resided in monopolin subunits (Mam1, Csm1, Hrr25 and Lrs4), Spo13^MOKIR^ or the spindle midzone protein Ase1^PRC1^, since this was also identified in the transposon-based screen (Rabitsch et al., 2003). Isolates that failed the complementation test were subjected to bulk segregation analysis followed by whole genome sequencing (Table S1; Materials and Methods). Together, this revealed that multiple isolates carried mutations in monopolin subunits including Mam1 (37%), Csm1 (14%) and Hrr25 (11%). Curiously, none of the isolates carried mutations in the fourth monopolin subunit Lrs4. We also identified several mutations in Spo13^MOKIR^ and Ase1^PRC1^. The remaining 36 isolates had causal mutations in 21 different genes (Figure 1C; Table S1). We confirmed that the 15 most confident candidate mutations identified bioinformatically were causal by *de novo* mutagenesis (Figure 1D). This revealed that, in addition to monopolin and MOKIR, mutations in genes encoding proteins involved in spindle pole body duplication (*SFI1*, *CDC55*), spindle midzone stability (*ASE1*), kinetochore function (*ASK1*, *SPC34*, *SPC105*), a cyclin (*CLB4*) and several kinases (*CDC7*, *CDC5* and *KNS1*) rescue the spore viability of *spo11Δ spo12Δ* cells. Interestingly, variant calling after whole genome sequencing identified high penetrance mutations in genes encoding both the chromosome passenger complex subunit *SL15^INCENP^* and the spindle checkpoint protein *MAD2*. During bulk segregation analysis, rescue of *spo11Δ spo12Δ* occurred at a 1:3 ratio for this isolate, suggesting two mutations acting in combination, which we confirmed by de novo mutant generation (Figure 1D). Therefore, the screen is robust and has the capacity to identify genes important for chromosome segregation and cell cycle regulation. Importantly, although several mutations occur in essential genes with known roles in mitosis, their recovery in the screen indicates that they support mitotic growth. We therefore conclude that effective meiosis I segregation places an increased dependence on diverse chromosome segregation pathways.

### Mutations in monopolin and Spo13^MOKIR^ define functional interfaces important for mono-orientation

The position and type of mutation can inform on protein function. We therefore directly sequenced the relevant monopolin genes, as well as *SPO13* and *ASE1*, in isolates where these genes were identified in the complementation analysis (Figure 2A; Table S1). Of the 14 *mam1* isolates sequenced, 7 carried frameshift or truncation mutations, while the remaining mutations occurred in regions that interact with Hrr25^CK1^ (3) or Csm1 (1) (Figure 2B and C). Due to its roles in multiple cellular processes, Hrr25^CK1^ is the only essential monopolin subunit (Petronczki et al., 2006). Accordingly, the 8 *hrr25* isolates sequenced carried either G12D (4) or R294C/H (4) mutations. Hrr25 G12 is a key residue for Mam1 interaction, while R294 lies just outside the characterised Mam1 interaction domain (Ye et al., 2016) and both residues were previously identified in a targeted screen to identify mono-orientation-defective alleles (Petronczki et al., 2006). All 13 *csm1* mutations sequenced mapped to the Mam1-binding interface, and represented 10 distinct mutations, underscoring the importance of the extended interface (Figure 2C). Curiously, no mutations in Lrs4, or the Csm1-Lrs4 interface were identified in our analyses. Because Csm1-Lrs4 also independently plays roles in rDNA organisation, mutations that disrupt this complex may cause growth defects that prevent recovery in the screen. Of the 5 *spo13* isolates sequenced, 1 carried a truncation mutation, 1 a start site loss mutation and 3 carried mutations in the Polo box domain (PBD) binding region, which is important for mono-orientation (Matos et al., 2008; Galander et al., 2019). Therefore, the screen was highly effective in the identification of functional interfaces that are important for mono-orientation.

**Figure 2.**
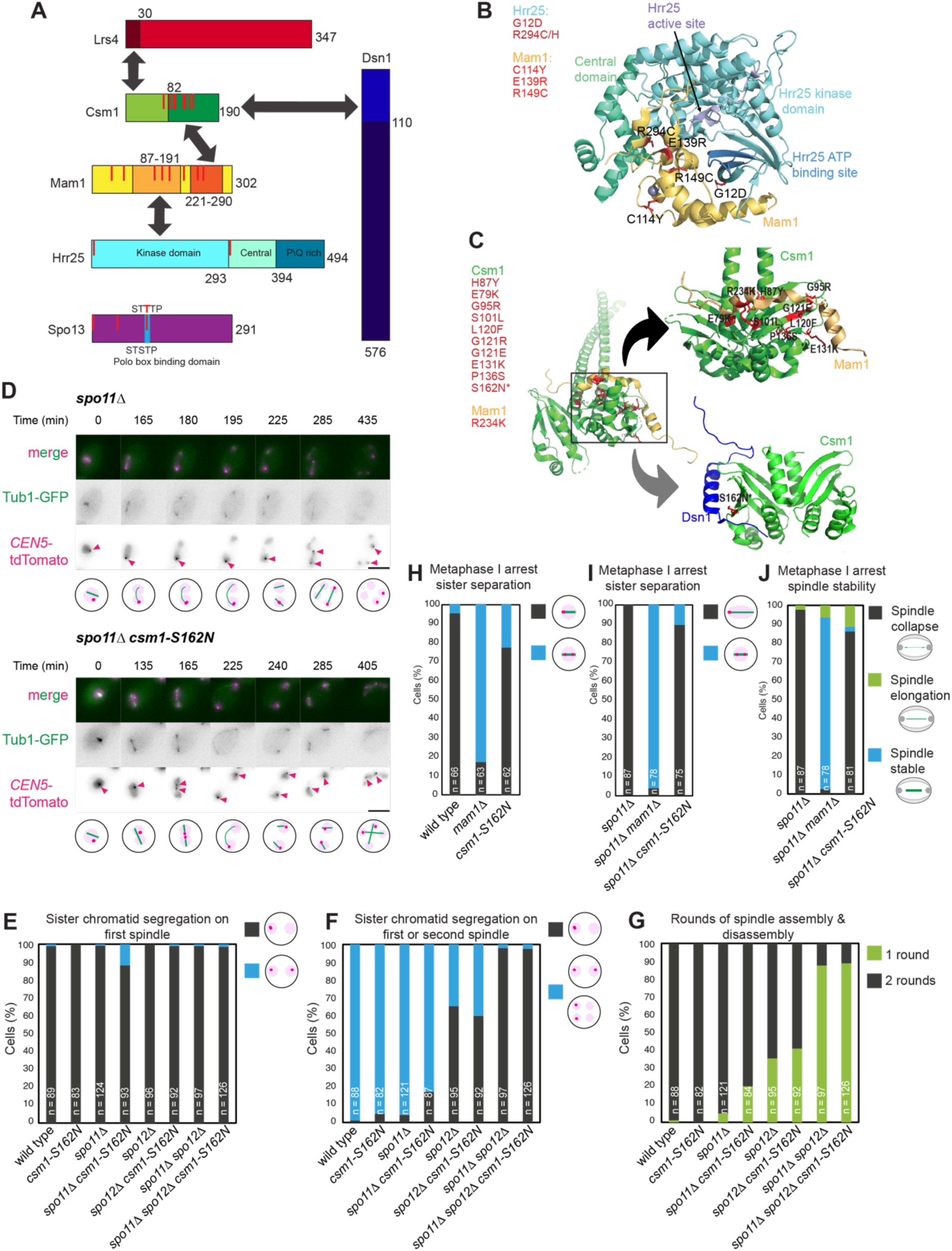
Mutations in monopolin disrupt key interfaces. (A) Schematic of monopolin subunits (Lrs4, Mam1, Csm1, Hrr25), together with the kinetochore receptor Dsn1, and the MOKIR, Spo13, showing the domains of interaction (arrows) and the position of mutations identified in the screen (red vertical lines). (B and C) Modelling of mutations identified in the screen onto previously determined structures of Hrr25-Mam1 (B), Csm1-Mam1 and Csm1-Dsn1 (C) showing that missense mutations cluster at these key interfaces. Structures shown are Csm1-Dsn1 (6MJB; (Plowman et al., 2019)), Hrr25-Mam1 (5CYZ; (Ye et al., 2016)) and Csm1-Mam1 (5KTB; (Corbett and Harrison, 2016). *Note that the Csm1-Dsn1 structure is from *Candida glabrata* proteins where S163 corresponds to residue S161 in *S. cerevisiae*. (D-G) A Csm1 mutation predicted to disrupt its interaction with kinetochore receptor Dsn1 impairs co-segregation of sister chromatids in meiosis I. Live imaging of cells carrying *CEN5-tdTomato* and *TUB1-GFP*, which label a single chromosome (magenta) or the spindle (green) respectively, undergoing meiosis. (D) Representative images of *spo11Δ* and *spo11Δ csm1-S162N* cells, showing correct co-segregation and splitting of *CEN5-tdTomato* at meiosis I, respectively. Magenta arrowheads indicate *CEN5-tdTomato* foci. Schematics below depict images above. (E and F) The percentage of cells of the indicated strains that separate *CEN5-tdTomato* foci either on the first (E) and/or second (F) spindle. (G) The percentage of cells that undergo one or two rounds of spindle assembly/disassembly is given for the indicated strains. In E-G, the number of cells scored is indicated. (H-J) Increased sister kinetochore bi-orientation at metaphase I in *csm1-S162N* cells. Live imaging of *CEN5-tdTomato TUB1-GFP* cells carrying *pCLB2-CDC20* to induce a metaphase I arrest. (H and I) The percentage of cells where split *CEN5-tdTomato* foci were observed was scored for the indicated genotypes. (J) The behaviour of the metaphase I spindle in the *spo11Δ* background was scored for the indicated genotypes, with a stable or elongated spindle being characteristic of bi-orientation rather than mono-orientation.

For reasons that are unclear, the complementation test failed to identify monopolin mutations in some isolates where they were likely causal. Notably, whole genome sequencing and *de novo* mutation confirmed causality of *csm1-S162N* in rescue of *spo11Δ spo12Δ* spore viability (Figure 1D). Interestingly, this residue lies in the Csm1 interface that binds the N terminus of the kinetochore protein Dsn1 and adjacent residue L161 is important for Dsn1 and Mif2 binding in vitro (Corbett et al., 2010) (Figure 2A). We therefore tested the effect of *csm1-S162N* on sister chromatid co-segregation *in vivo* in a clean diploid background. Cells carrying heterozygous tdTomato labels on the centromere of chromosome V (*CEN5-tdTomato*) and Tub1-GFP to label the spindle were induced to undergo meiosis and live imaged. Since chiasma can obscure mono-orientation defects, we also carried out this experiment in the *spo11Δ* background (Figure 2D-G). In wild-type and *spo11Δ* cells, as expected, sister *CEN5-tdTomato* foci co-segregated in meiosis I, but split only in meiosis II (Figure 2D-F). While *csm1-S162N* cells also accurately co-segregated sister chromatids during meiosis I, around 11% of *spo11Δ csm1-S162N* cells split sister chromatids to opposite poles (Figure 2D and E). Consistently, *csm1-S162N* reduced the proportion of cells that underwent two rounds of spindle assembly in the *spo11Δ* background (Figure 2F). Unexpectedly, *spo11Δ spo12Δ csm1-S162N* cells did not split sister chromatids in meiosis I (Figure 2D and E). While the underlying reasons remain unexplained, we speculate that the requirement to establish tension-generating kinetochore-microtubule attachments is relaxed in *spo12Δ.* To confirm that *csm1-S162N* impairs sister kinetochore mono-orientation, rather than uncoupling chromosome segregation from spindle assembly (Figure 1A), we arrested cells in metaphase I by depleting Cdc20 and scored the splitting of sister *CENV–tdTomato* foci (Figure 2H-J). In wild-type cells, *CENV-tdTomato* foci rarely split, consistent with robust mono-orientation, whereas over 80% of *mam1Δ* cells exhibited separation, indicating loss of mono-orientation. In contrast, ∼20% of *csm1-S162N* cells showed split *CENV-tdTomato* foci, consistent with a partial defect (Figure 2H). Similar results were observed in a *spo11Δ* background (Figure 2I). As a further test, we examined spindle morphology in the *spo11Δ* background, where the absence of chiasmata leads to premature spindle elongation and collapse due to lack of tension (for examples of spindle phenotypes see Figure 3A). In contrast, loss of mono-orientation, as in *mam1Δ*, allows sister kinetochore biorientation so that cohesin generates tension, producing short stable spindles (Figure 2I). *csm1-S162N* also increased spindle stability of *spo11Δ* cells, albeit modestly (Figure 2J). Together, these data indicate that *csm1-S162N* modestly impairs mono-orientation, as revealed during metaphase I arrest, but that this defect is compensated during meiotic progression, in a manner dependent on chiasmata/Spo11. Consistent with its modest effects on mono-orientation when cells were allowed to progress through meiosis, *csm1-S162N* had little effect on spore viability in an otherwise wild-type background (Figure S2A and B). We conclude that the Dsn1-binding region of Csm1 is important for robust mono-orientation *in vivo* and that backup mechanisms, including chiasmata, can rescue segregation when mono-orientation is modestly impaired. Overall, our EMS-based mutagenesis screen is a powerful approach to define functionally important protein surfaces involved in meiosis I chromosome segregation.

**Figure 3.**
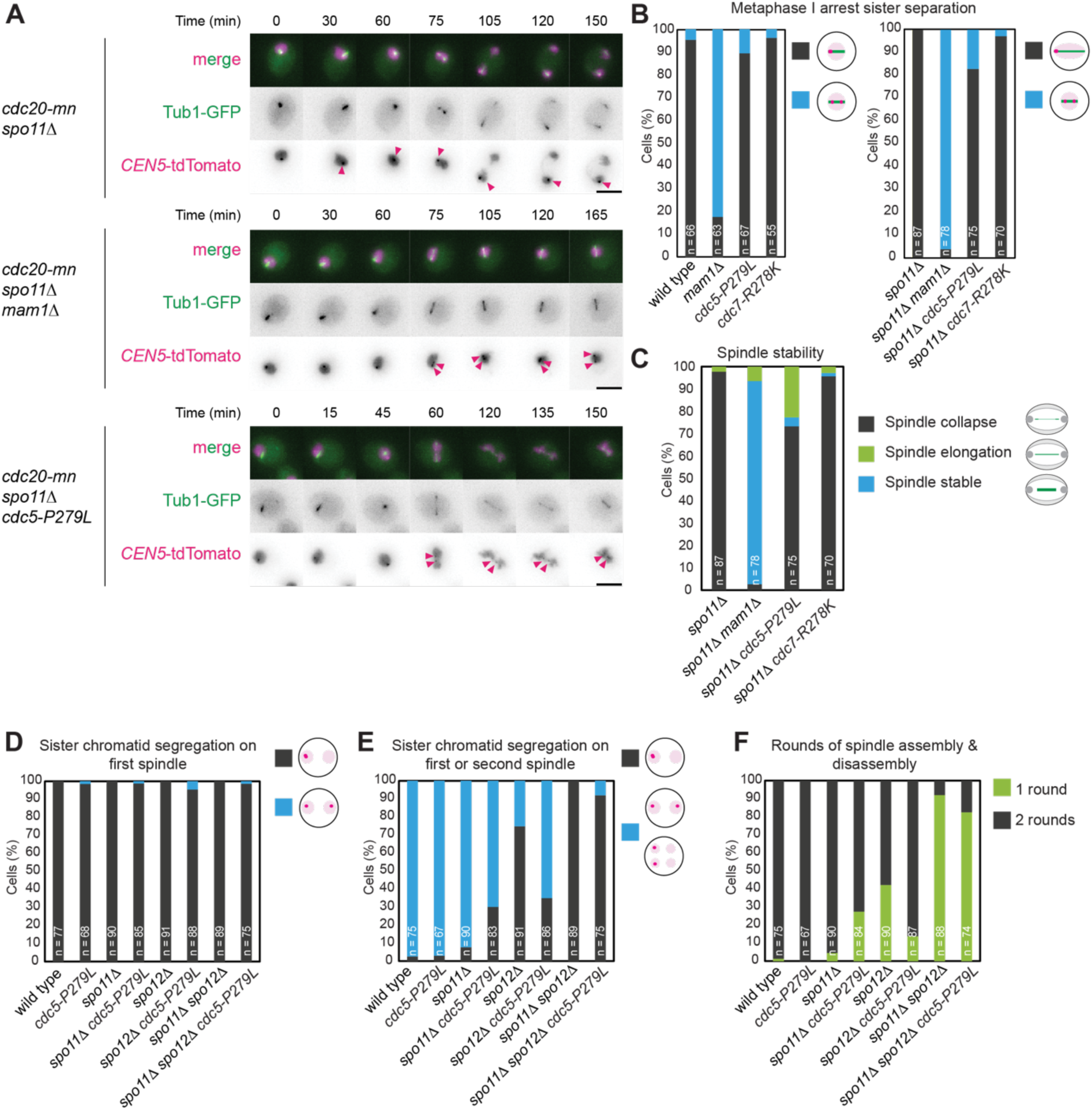
Meiotic kinases promote mono-orientation and co-ordination. (A-C) *cdc5-P279L*, but not *cdc7-R278K* shows defective mono-orientation. Live imaging of metaphase I-arrested cells as in Figure 2H-J. Representative images (A) and the percentages of cells with split *CEN5-tdTomato* foci (B) or the indicated spindle morphologies (C) are shown. (D-F) Sister chromatids typically co-segregate in *cdc5-P279L* meiosis I cells, however, *cdc5-P279L* increases the proportion of *spo12Δ* cells undergoing a sister chromatid segregation on a second spindle. Live imaging of cells undergoing meiosis as in Figure 2D-G, with the percentage of cells that separate *CEN5-tdTomato* foci on the first (D) and/or second (E) spindle, together with the percentage of cells undergoing one or two rounds of spindle assembly/disassembly (F) are shown.

### Meiotic kinases implicated in mono-orientation also promote meiosis I segregation through additional mechanisms

Our screen identified mutant alleles (*cdc5-P279L* and *cdc7-R278K*) of two kinases required for mono-orientation, Cdc5 and Cdc7 (Matos et al., 2008; Lee and Amon, 2003; Clyne et al., 2003). *cdc5-P279L* rescued *spo11Δ spo12Δ* spore viability to a similar extent to *mam1*1, while *cdc7-R278K* showed a more moderate rescue (Figure 1D). Both kinases are essential for viability, suggesting that the identified mutations have only modest or meiosis-specific effects. Indeed, in an otherwise wild-type background, *cdc5-P279L* modestly reduced spore viability, whereas *cdc7-R278K* had no detectable effect (Figure S2). To determine whether these mutations impair mono-orientation, we scored separation of heterozygous *CEN5-tdTomato* foci in metaphase I-arrested cells (by depletion of Cdc20), either in wild-type or *spo11Δ* backgrounds. *cdc5-P279L* cells showed a low frequency of *CEN5-tdTomato* separation and corresponding increase in spindle stability in the *spo11Δ* background, indicating a modest defect in mono-orientation (Figure 3A-C). In contrast, *cdc7-R278K* was proficient in mono-orientation, indicating rescue of *spo11Δ spo12Δ* spore viability through a different mechanism (see below). However, in contrast to metaphase I cells, *cdc5-P279L* cells only rarely split heterozygous *CEN5-tdTomato* foci on the first spindle when allowed to progress through meiosis, even in the *spo11Δ* background (Figure 3D). This indicates that the weak mono-orientation defects at metaphase I in *cdc5-P279L* cells are somehow compensated for during anaphase I. Interestingly, *cdc5-P279L* increased the fraction of *spo12Δ*, and to a lesser extent *spo11Δ spo12Δ*, cells that underwent a second round of spindle assembly and disassembly, accompanied by increased segregation of sister chromatids on this second spindle (Figure 3E and F). Together, these results suggest that *cdc5-P279L* affects both mono-orientation and the control of chromosome segregation after meiosis I, indicating that this allele compromises multiple functions of Cdc5 in the meiotic division program.

A similar but weaker phenotype was observed for *cdc7-R278K*, which increased the proportion of *spo12Δ* cells undergoing a second round of spindle assembly in which sister chromatids segregate (Figure S3A-C). In both *cdc5-P279L* and *cdc7-R278K*, this effect was dampened by *spo11Δ*, which reduced the proportion of cells that underwent a second round of spindle assembly (Figure 3F and Figure S3C). This is consistent with the idea that, in the absence of chiasmata, chromosomes more readily clear the spindle midzone, reducing the likelihood that a second round of segregation will occur (Marston et al., 2003; Bizzari and Marston, 2011). It is, nevertheless, surprising that *cdc5-P279L* and *cdc7-R278K* show very little sister chromatid separation in the *spo11Δ spo12Δ* background used for the screen (Figure 3E and Figure S3A and B). One possible explanation is that the screen was performed in haploids, whereas live imaging was conducted in larger diploid cells, which may allow chromosomes to clear the spindle midzone more readily.

We also identified two truncation mutants *(kns1-W692X and kns1-W604X)* of the kinase Kns1 in the screen and confirmed it to be causal (Figure 1D and Table S1). However, the mechanism by which this mutation rescues *spo11Δ spo12Δ* spore viability remains unclear, as live imaging of *kns1-Δ604-738* mutants revealed no obvious chromosome segregation phenotype (Figure S3D-F). A previous study also reported no sporulation defects in *kns1Δ* cells (Padmanabha et al., 1991), therefore the basis for the isolation of *kns1* mutants in the screen remains unexplained.

Together, these results indicate that kinase mutations recovered in the screen can influence meiosis I chromosome segregation through multiple mechanisms, including mono-orientation as well as altered control of chromosome segregation after meiosis I. Importantly, the screen also identifies mutations in essential genes, providing insight into how their functions are adapted for meiosis.

### A weak spindle midzone prevents meiosis I chromosome segregation

We recovered 16 isolates carrying mutations in *ASE1*, encoding a conserved spindle midzone crosslinker, and representing at least 12 distinct alleles (Table S1). *ASE1* was also identified in the earlier transposon-based screen (Rabitsch et al., 2003). In our isolates, mutations were distributed throughout the protein, suggesting that rescue of *spo11Δ spo12Δ* spore viability is due to loss of Ase1 function. We hypothesised that *ASE1* belongs to the class of identified genes which are particularly important for meiosis I chromosome segregation (Figure 1A). To test this idea, we generated an Ase1 truncation mutant, mimicking one of the screen isolates, *ase1-Δ473-886*. The newly generated *ase1-Δ473-886* mutant in a clean background, rescued the spore viability of *spo11Δ spo12Δ* to a similar extent to the isolate from the screen (Figure 4A). In an otherwise wild-type background, the spore viability of *ase1-Δ473-886* was modestly reduced (Figure S2B). Despite this, live imaging revealed that *ase1-Δ473-886* diploid cells carrying heterozygous *CEN5-tdTomato* and *TUB1-GFP* could complete two meiotic divisions with sister chromatids segregating only after the second division and this was unchanged in the *spo11Δ* background (Figure 4B-D). However, *ase1-Δ473-886* increased the fraction of *spo12Δ* cells that assembled a second spindle on which sister *CEN5-tdTomato* foci segregate to opposite poles (Figure 4E). As a result, *spo11Δ spo12Δ ase1-Δ473-886* cells ultimately produce dyads with an even distribution of sister chromatids, explaining their viability and recovery in the screen. Thus, defective Ase1 causes abortive meiosis I chromosome segregation in the *spo12Δ* background. Instead, sister chromatids segregate at the time of meiosis II, when monopolin and arm cohesion have been lost and sister chromatids biorient. We conclude that Ase1 is crucial to ensure proper completion of meiosis I segregation to avoid de-coupling of the chromosome segregation programme and the meiotic spindle cycle.

**Figure 4.**
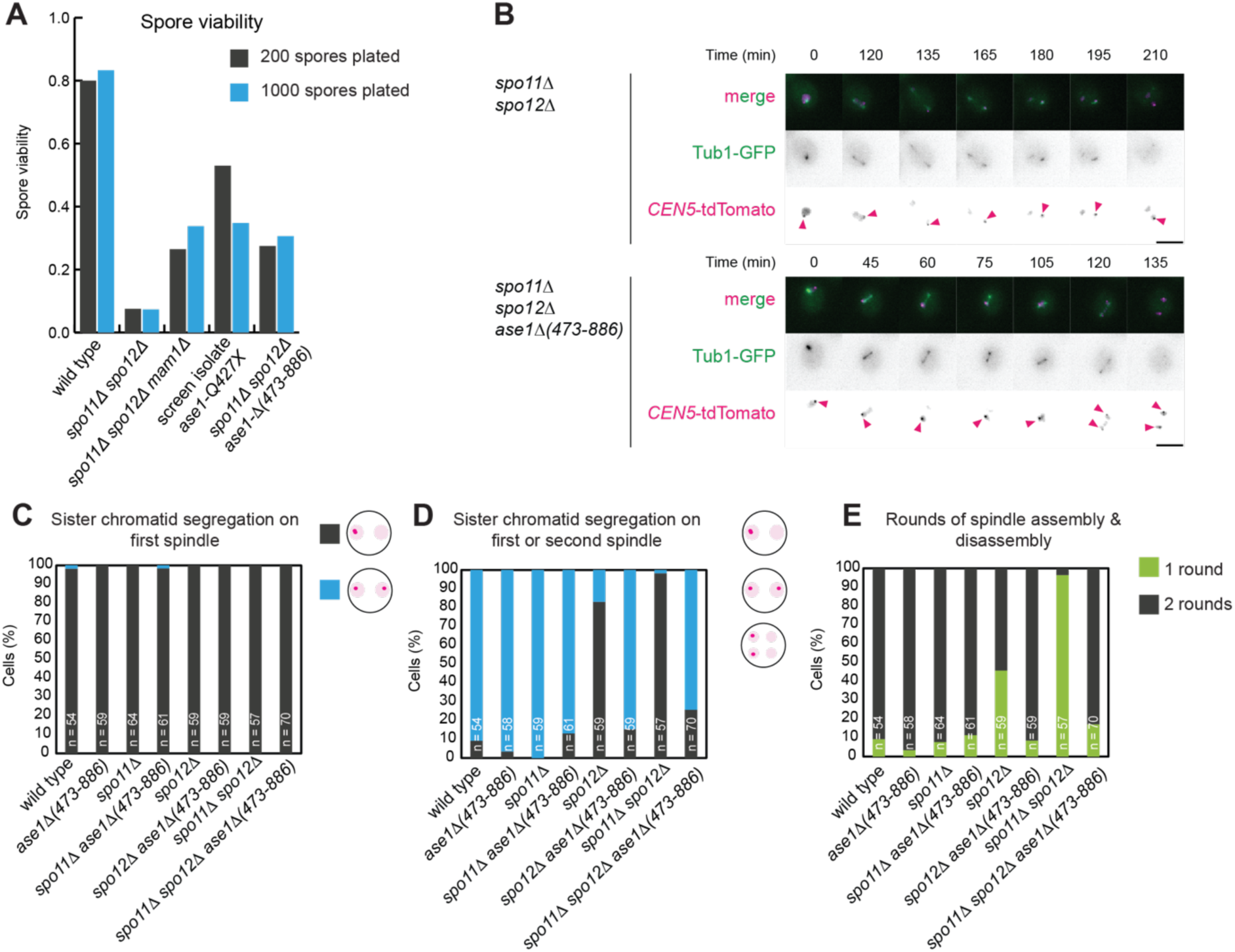
Coordinated and sequential meiotic divisions require spindle midzone stability. (A) Comparison of *spo11Δ spo12Δ* rescue in the screen isolate with the *de novo* generated mutation for *ase1-Δ* (473-886). (B-E) *ase1-Δ (473-886)* increases the frequency of second spindle formation on which sister chromatids segregate. Live imaging of cells carrying *CEN5-tdTomato* and *TUB1-GFP* as in Figure 2D-G. (B) Representative images, (C and D) the percentage of cells in which sister chromatids segregate on the first (C) and/or second (D) spindle. (E) The percentage of cells undergoing one or two rounds of spindle assembly/disassembly.

### Delayed spindle pole body separation results in omission of meiosis I chromosome segregation

We hypothesised that an alternative mechanism by which the chromosome segregation programme could become uncoupled from the meiotic spindle cycle would be a delay in spindle pole body duplication (Figure 1A). Consistent with this idea, the screen recovered mutations in two conserved genes involved in spindle pole body separation, *SFI1* and *CDC55* (Table S1). Sfi1 is a component of the bridge structure important for spindle pole body duplication (Rüthnick and Schiebel, 2016), while Cdc55 is a regulatory subunit of the protein phosphatase 2A (PP2A) (Nilsson, 2018). Interestingly, the three *sfi1* mutations recovered in the screen (*sfi1-R788L*, *sfi1-P817S* and *sfi1-P856S*) all lie in Cdk1 consensus sites (Elserafy et al., 2014). We also identified 4 distinct mutations in *CDC55* (Table S1) and based on high conservation focused on *CDC55-G43D*. In an otherwise wild-type background, spore viability was modestly reduced in *sfi1-R788K* and more strongly affected in *cdc55-G43D* (Figure S2). Despite undergoing only one round of spindle formation, *spo11Δ spo12Δ cdc55-G43D* cells segregated *CEN5-tdTomato* to opposite poles as soon as the spindle formed (Figure S4A-C). Unlike in *mam1Δ* cells where centromeric cohesion prevents elongation of the first spindle (Toth et al., 2000), *CEN5-tdTomato* foci separated immediately upon spindle formation, suggesting that the single spindle in *spo11Δ spo12Δ cdc55-G43D* forms at the time of meiosis II when centromeric cohesion has already been lost. To further test this, we imaged spindle pole bodies (Spc42-tdTomato) together with meiotic cohesin (Rec8-GFP). In wild type, *spo11Δ*, *spo12Δ* and *spo11Δ spo12Δ* cells, two SPB foci are invariably observed prior to bulk Rec8 loss. In contrast, in the *cdc55-G43D* background, bulk Rec8 was lost prior to SPB separation, at which time only the minor centromeric Rec8 pool remained (Figure 5A and B). Consistent with its ability to support mitotic growth, *cdc55-G43D* supported a single round of SPB duplication/separation in the majority (∼80%) of cells, which was greatly delayed (Figure 5C and D). This suggests that *cdc55-G43D* disrupts a regulatory mechanism that permits two consecutive SPB duplication/separation events in meiosis, rather than being generally required for SPB duplication and separation. A similar, albeit less severe, phenotype was observed in *sfi1-R788K* cells, where SPB separation occurred after bulk Rec8 loss in a fraction of cells, but was nevertheless significantly delayed (Figure 5E-H). Interestingly, *spo11Δ* increased the fraction of *sfi1-R788K* cells where SPB separation occurred after bulk Rec8 loss and extended the time between these events (Figure 5E-I). Potentially the absence of chiasmata facilitates occupancy of all kinetochores by microtubules emanating from monopolar spindles, thereby more readily satisfying the spindle assembly checkpoint resulting in earlier Rec8 loss. Therefore, by delaying SPB separation, *cdc55-G43D* and *sfi1-R788K* uncouple the timing of spindle formation from the chromosome cycle. As a result, the first spindle forms at the time of meiosis II, when chromosomes are in a state that supports sister kinetochore biorientation (Figure 5I). *SFI1* is an essential gene and *CDC55* null mutations have pleiotropic effects on mitosis and meiosis; however, the alleles recovered in our screen support vegetative growth, indicating that modest perturbations to these proteins are tolerated during mitosis (Anderson et al., 2007) but disrupt the timing of spindle pole body duplication during meiosis. We conclude that meiosis I spindle poles are highly sensitive to perturbations that disrupt duplication timing, risking uncoupling of chromosome segregation from the meiotic spindle cycle.

**Figure 5.**
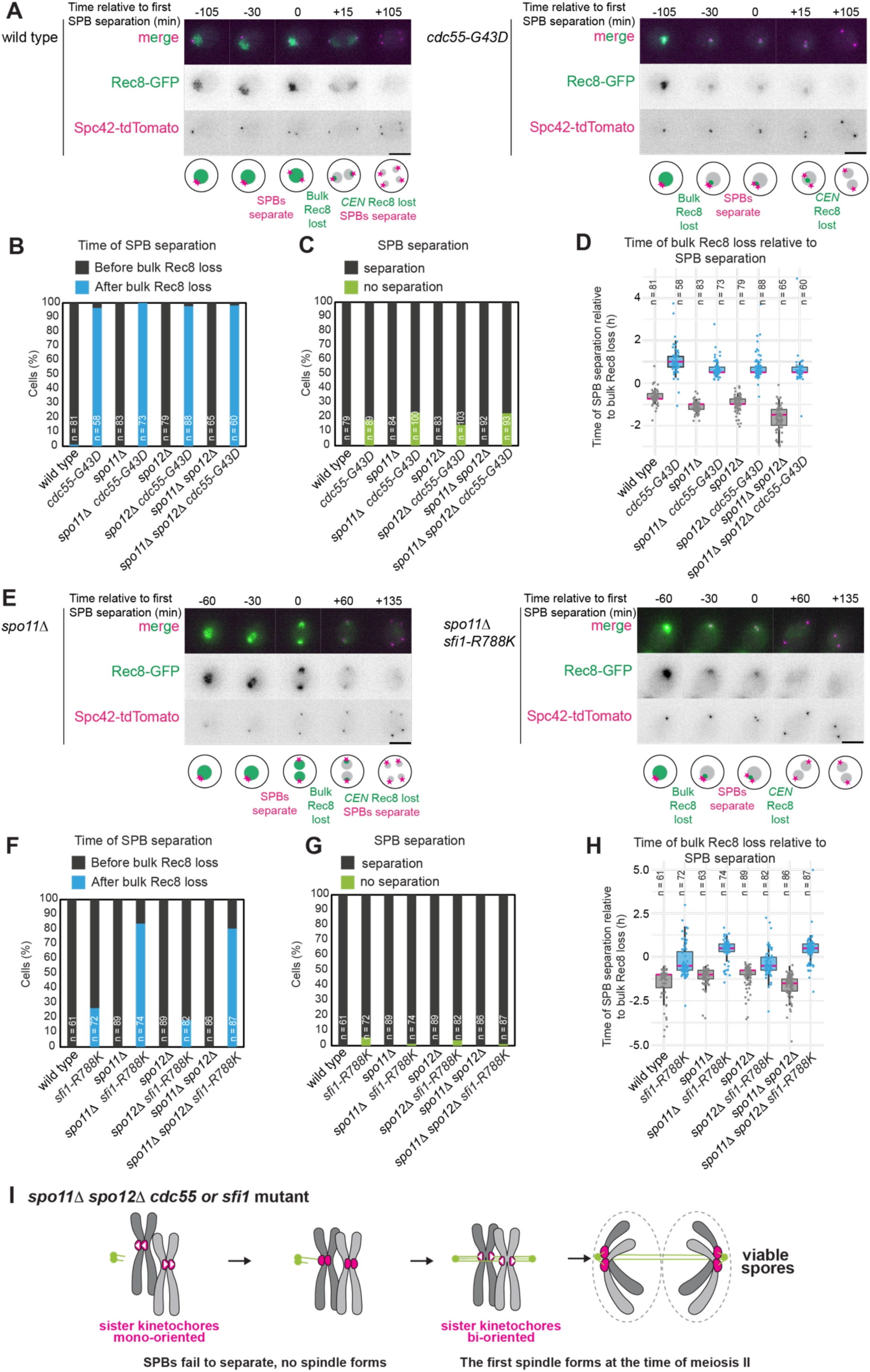
Timely spindle pole body separation is critical for homolog segregation in meiosis I. (A-D) SPB separation occurs only after the first round of cohesin loss in *cdc55-G43D* cells. Live imaging of cells carrying *REC8-GFP* (cohesin) and *SPC42-tdTomato* (SPBs) and induced to undergo meiosis. (A) Representative images, and illustrative schematics below, of wild type and *cdc55-G43D* cells. (B) The percentage of cells where bulk Rec8 loss occurred before or after SPB duplication was scored for the indicated genotypes and number of cells. (C) The percentage of cells undergoing at least one round of SPB separation was scored. Note that these cells still lose bulk Rec8. (D) The time interval between bulk Rec8 loss and SPB separation was determined for the indicated number of cells and genotypes. (E-I) *sfi1-R788K* delays SPB duplication, with the greatest effect in the *spo11Δ* background. Live imaging as in A-D. (E) Representative images. (F) Percentage of cells where Rec8 was lost prior to SPB separation. (G) Percentage of cells undergoing at least one round of SPB separation overall. (H) Time between bulk Rec8 loss and SPB separation. (I) Schematic showing consequences of failed SPB duplication prior to meiosis I. Lack of spindle formation in meiosis I does not prevent onset of meiosis II cell cycle state, such that mono-orientation is lost prior to spindle assembly and sister kinetochores bi-orient and segregate.

### Meiosis I is highly sensitive to outer kinetochore defects

Outer kinetochore proteins represented another class of mutations identified in the screen (Table S1). These included mutations in the Dam1 complex (*spc34-G81E* and *ask1-P217L*) and in the KMN complex (*spc105-R102C*). We reasoned that by weakening kinetochore–microtubule attachments, these mutations could impair chromosome movement during meiosis I in the *spo12Δ* background, resulting in an abortive meiosis I division similar to that observed in *ase1* mutants and ultimately leading to segregation of sister chromatids on the second meiotic spindle (Figure 1A). Consistent with this idea, all three outer kinetochore mutants exhibited this phenotype, at least in the *spo12Δ* background (Figure 6A-D; Figure S5). In the *ask1-P217L* and *spc105-R102C*, but not *spc34-G81E* mutants, the proportion of *spo12Δ* cells undergoing two rounds of spindle assembly and disassembly was also modestly increased (Figure 6D, Figure S5C and F). Rather than affecting spindle stability as in *ase1-Δ473-886*, it is likely that *spc34-G81E*, *ask1-P217L* and *spc105-R105C* impair the attachment of chromosomes to microtubules. As a result, chromosomes would fail to segregate efficiently at meiosis I, increasing the likelihood that they are eventually segregated on the meiosis II spindle. Importantly, the *spo12Δ* background provides a sensitised system for these defects to be revealed, since *spc34-G81E*, *ask1-P217L* and *spc105-R105C* do not greatly compromise spore viability in an otherwise wild-type background (Figure S2).

**Figure 6.**
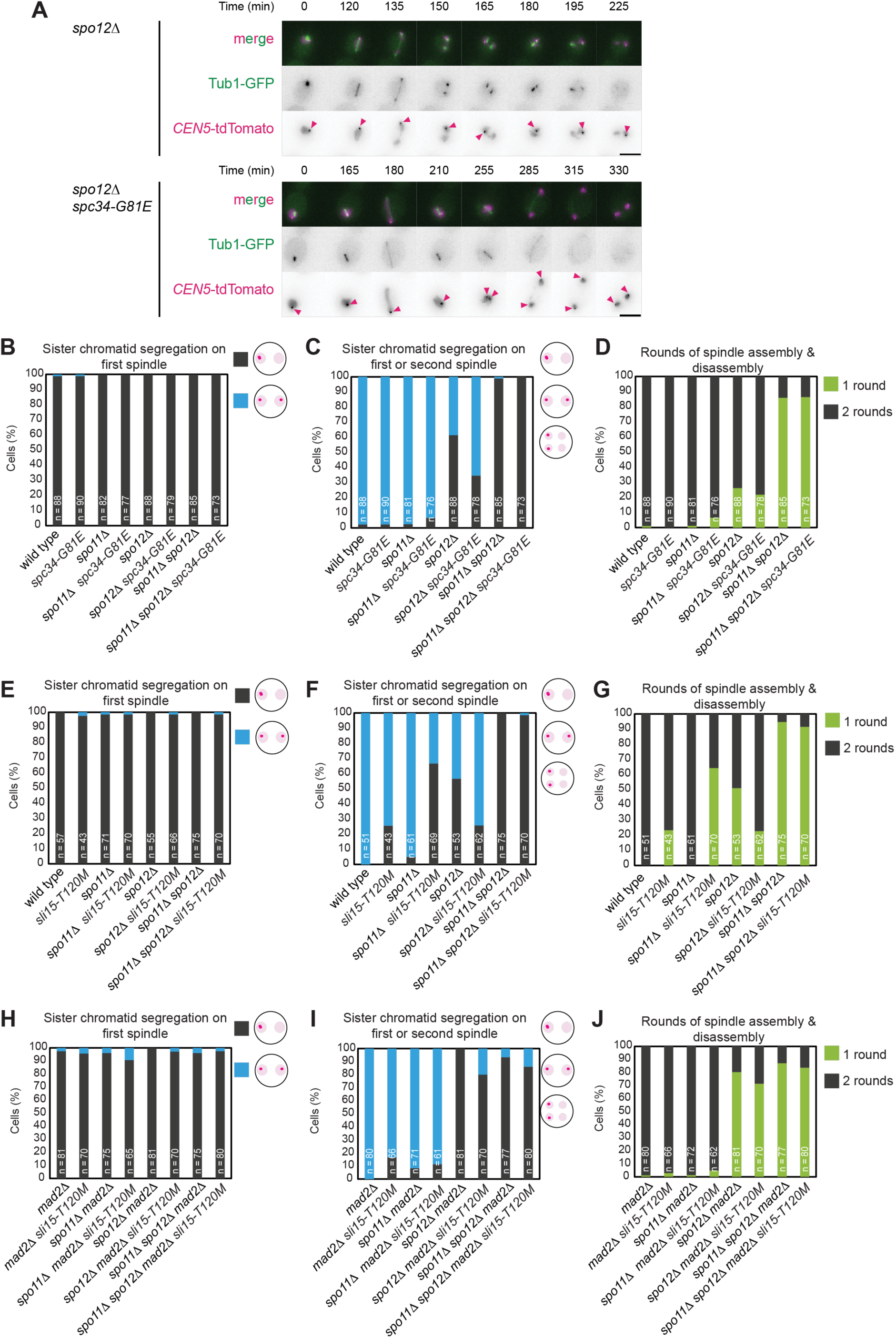
Kinetochore defects delay chromosome segregation until meiosis II. (A-D) *spc34-G81E* modestly increases the fraction of *spo12Δ* cells that segregate sister chromatids on the second spindle, without increasing the frequency of second spindle formation. Live imaging as in Figure 2D-G with representative images (A), the percentage of cells separating *CEN5-tdTomato* foci on the first (B) and/or second (C) spindle, together with the percentage of cells undergoing one or two rounds of spindle assembly/disassembly (D) are shown. (E-J) *sli15-T120M* and *mad2Δ* in combination increase the frequency of *spo12Δ* cells that separate *CEN5-tdTomato* foci on a second spindle. Live imaging as in Figure 2D-G with the percentage of cells separating *CEN5-tdTomato* foci on the first (E and H) and/or second (F and I) spindle, together with the percentage of cells undergoing one or two rounds of spindle assembly/disassembly (G and J) are shown for the indicated genotypes.

Interestingly, a further isolate identified in the screen also appeared to rescue *spo11Δ spo12Δ* spore lethality potentially through a kinetochore-related mechanism. During backcrossing and bulk segregant analysis, we noticed that only ∼25% of segregants exhibited the rescue phenotype. This deviates from the 50% phenotype penetrance expected for a single-gene mutation, suggesting that the rescue instead arises from mutations in two independently segregating genes. Consistently, whole genome sequencing revealed almost 100% penetrance with both *SLI15* and *MAD2* mutations (*sli15-T120M* and *mad2-3QX*). Because *MAD2* was truncated early in its coding sequence, we generated *sli15-T120M* and introduced it into a *mad2Δ* strain. The double *sli15-T120M mad2Δ* mutant showed the strongest rescue of *spo11Δ spo12Δ* viability of all isolates tested, while the single mutants provided little to no rescue (Figure 1D). In an otherwise wild-type background, *sli15-T120M mad2Δ* greatly reduced spore viability. In comparison, *mad2Δ* alone had a moderate effect, as expected (Mukherjee et al., 2024), while *sli15-T120M* alone had only a minor effect on spore viability (Figure S2). Live imaging further revealed that in the *spo12Δ* background, the frequency of sister chromatid segregation on the second spindle, and the proportion of cells undergoing two rounds of spindle assembly and disassembly, were modestly increased in the *sli15-T120M mad2Δ* mutant. A smaller increase was also observed in *sli15-T120M* alone, while *mad2Δ* alone had no detectable effect (Figure 6E-J).

Together, these findings indicate that defects in the spindle midzone, spindle pole bodies, or kinetochores represent three distinct routes by which chromosome segregation can become uncoupled from the meiotic spindle cycle, disrupting the normally strict order of homolog segregation in meiosis I followed by sister chromatid segregation in meiosis II.

## Discussion

Successful gamete formation during meiosis requires the coordinated action of multiple mechanisms that ensure the orderly execution of two sequential and distinct divisions. In particular, the switch in kinetochore orientation between meiosis I and II must be coupled to two rounds of spindle assembly and disassembly. By identifying mutants that restore sister chromatid segregation in *spo11Δ spo12Δ* cells, we show that this coupling depends on diverse components of the chromosome segregation machinery, including monopolin, meiotic kinases, spindle midzone factors, spindle pole body regulators and outer kinetochore proteins (Figure 7). Together, these findings define multiple pathways that contribute to this coordination and show how its disruption can lead to segregation errors.

**Figure 7.**
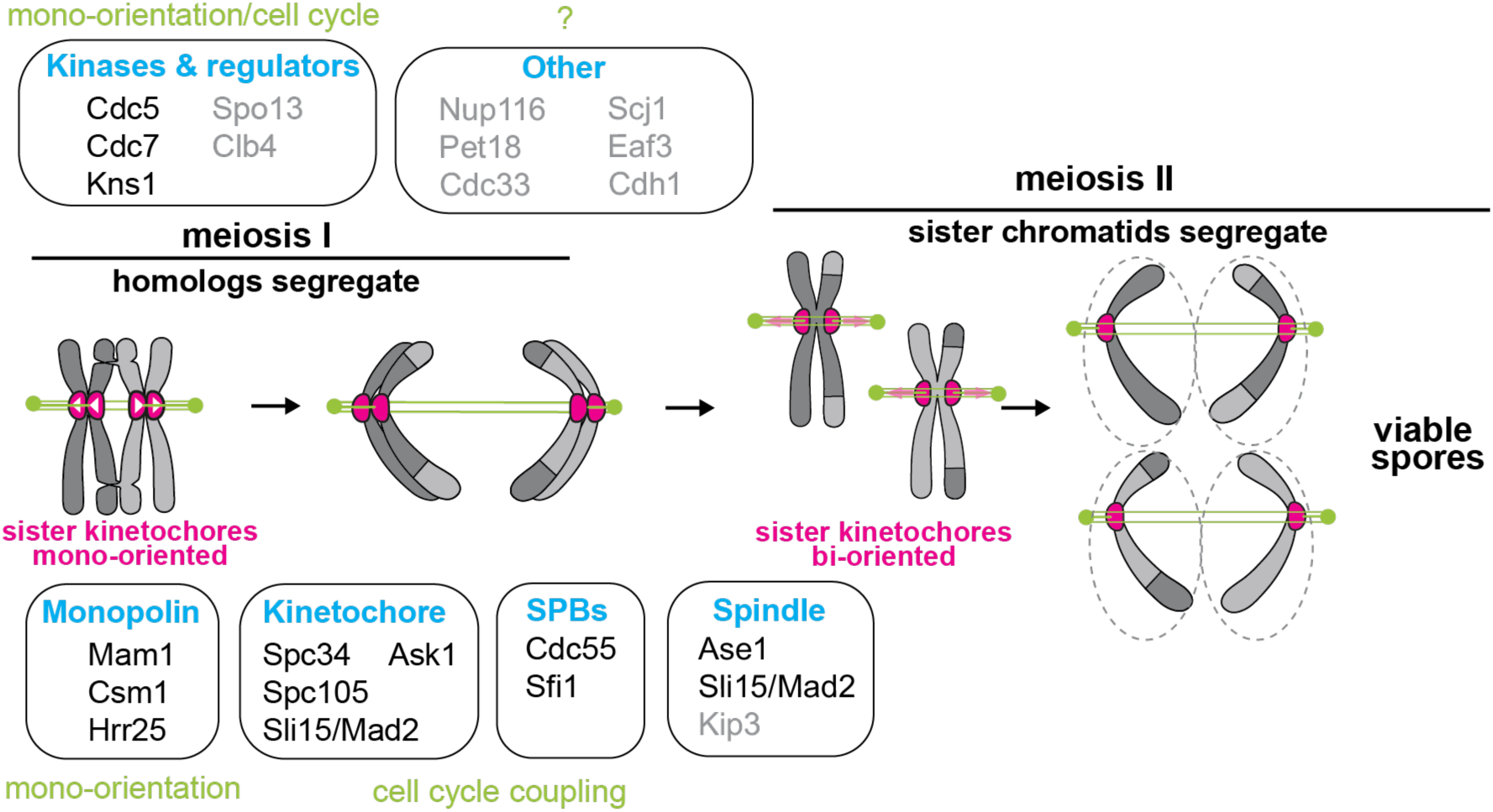
Meiosis I is highly sensitive to a range of segregation defects, leading to loss of coordination between meiotic divisions. Schematic summarising the proteins found in this study to be important for coordinating meiosis, along with the processes/complexes they contribute to. Proteins in black were verified as causative and the mechanism of *spo11Δ spo12Δ* rescue was established. Proteins in grey were identified through high variant allele frequency scores in whole genome sequencing as likely causative but have not been verified.

### Screening for meiotic mutants

Discovery of the cellular machinery required for meiosis is particularly challenging. Because germ cells and gametes are scarce in mammals and suitable *in vitro* culture systems are lacking, simple model organisms play an essential role. Nevertheless, even studies in yeast, one of the simplest meiotic systems, present considerable barriers to discovery. While genetic screens in haploid yeast have been extremely powerful for understanding mitosis, meiotic entry requires diploid cells, complicating the recovery and analysis of recessive mutations in similar screens. Despite this, classical screens in haploid homothallic (mating-type switching) or pseudodiploid strains successfully identified sporulation and recombination mutants and mapped them by complementation (Nairz and Klein, 1997; Malone et al., 1991; Tsuboi, 1983; Esposito et al., 1970; Blyth et al., 2018). More systematic approaches, including candidate screens based on meiotically upregulated genes (Toth et al., 2000) and genome-wide deletion collections (Marston et al., 2004; Enyenihi and Saunders, 2003; Briza et al., 2002), have also identified many genes required for sporulation and meiosis. Chemical mutagenesis offers a complementary strategy that can uncover informative alleles, and the availability of whole-genome sequencing now makes it straightforward to identify causal mutations. However, this approach requires backcrossing to remove the large number of background mutations introduced by mutagenesis. Because meiosis requires expression of both mating-type programmes, subsequent mating, and therefore backcrossing, is prevented. To overcome this limitation, we engineered a conditional pseudodiploid system in which one mating-type programme can be selectively degraded. This allows mating competence to be restored after meiosis, enabling subsequent crosses and identification of causative mutations by whole genome-sequencing.

Meiotic defects, particularly those affecting chromosome segregation, produce inviable spores, preventing recovery of the mutant genotype. Several screens have therefore used genetic backgrounds in which alterations to the segregation programme restore, rather than eliminate, spore viability (Rabitsch et al., 2003; Yokobayashi and Watanabe, 2005; Malone et al., 1991). Similarly, we exploited the *spo11Δ spo12Δ* background to identify mutations that restore sister chromatid segregation and spore viability. The mutants we identified therefore provide insight into how kinetochore orientation is coupled to the meiotic spindle cycle.

### Mechanisms controlling mono-orientation in meiosis

The recovery of informative alleles affecting interfaces within the monopolin complex is a key outcome of our screen. In addition to frameshift or premature termination mutations in monopolin subunits and the MOKIR protein Spo13, we isolated mutations predicted to disrupt interactions between complex components, including the Csm1–Mam1 and Mam1–Hrr25 interfaces. Mutations in Spo13 mapped to the Polo-binding domain interaction motif. We also identified a mutation in Csm1 predicted, based on structural and in vitro studies (Plowman et al., 2019; Corbett and Harrison, 2016; Corbett et al., 2010), to disrupt interaction with the N-terminal region of the kinetochore protein Dsn1. Notably, we did not recover mutations within the Dsn1 N-terminus itself, despite this region being dispensable for viability in mitosis (Sarkar et al., 2013). Truncation or mutation of the Dsn1 N-terminus causes dominant mis-segregation in meiosis, providing a potential explanation for their absence in the screen (Sarkar et al., 2013; Plowman et al., 2019). Similarly, we did not identify mutations in the monopolin subunit, Lrs4, or in the region of Csm1 that interacts with Lrs4, suggesting that such mutations have dominant or otherwise deleterious effects on meiosis that preclude their recovery in the screen.

Similar screens have used the same principle of restoring balanced chromosome segregation in single-division meiosis to identify mono-orientation mutants. In *S. cerevisiae* this strategy led to the identification of monopolin subunits (Rabitsch et al., 2003), while in *S. pombe* it revealed the MOKIR protein Moa1 together with regulators of cohesin (Yokobayashi and Watanabe, 2005). A recent experimental evolution screen for suppressors of *spo11Δ* spore lethality isolated loss of function mutations in *CLB4*, including the *clb4-W246X* allele identified here, in addition to *SPO13^MOKIR^* and *MAM1* (Lumbroso et al., 2025). A conceptually similar screen in *Arabidopsis thaliana* identified cohesin subunits and their regulators, as well as the kinetochore protein CENP-C and a deSUMOylase as factors affecting mono-orientation (Singh et al., 2025). Notably, cohesin components or regulators were not recovered in our screen, despite our previous finding that Eco1-dependent acetylation of cohesin is required for both centromeric cohesion and mono-orientation in budding yeast meiosis (Barton et al., 2022). One possible explanation is that budding yeast possesses fewer cohesin variants than organisms such as *S. pombe* or plants, where multiple cohesin subunits provide greater functional redundancy. As a result, even modest perturbations of cohesin function in budding yeast may produce more severe meiotic defects, precluding recovery of such mutants.

### Coordination of chromosome segregation with the meiotic spindle cycle

Another key outcome from our screen is that the coordination of chromosome behaviour with the timing and organisation of the meiotic spindle can be disrupted in multiple ways. The mutants we identified define several distinct classes of defects, including perturbations to spindle midzone function, kinetochore–microtubule attachment, spindle pole body duplication and regulatory pathways. Our findings also show that the *spo11Δ spo12Δ* background provides a uniquely sensitive context in which to reveal such defects. In this background, when chromosome segregation during the first division is compromised, cells can undergo an additional round of spindle assembly along the same axis. It is thought that by constraining spindle pole body separation, failed chromosome segregation in meiosis I allows the two half spindles to meet, favouring a second round of spindle assembly on which those chromosomes can segregate (Marston et al., 2003; Bizzari and Marston, 2011; Mengoli et al., 2021). Mutations that impair chromosome movement, retention, or attachment during meiosis I therefore increase the likelihood of segregation during this later division. The diversity of mutant classes highlights that chromosome segregation can be uncoupled from the spindle cycle through distinct mechanisms. One class of mutants impairs chromosome coordination by disrupting meiosis I spindle assembly. This includes mutants of the spindle midzone protein Ase1, which fail to form a robust meiosis I spindle, as well as mutants of the SPB-separation factors Sfi1 and Cdc55, which do not produce a meiosis I spindle at all. Antiparallel dimerisation of Sfi1 forms a bridge between the new and old SPBs, a process regulated by Cdk1-dependent phosphorylation and Cdc14-dependent dephosphorylation (Avena et al., 2014; Elserafy et al., 2014). Strikingly, the *sfi1-R788K* mutation we identified lies in the C-terminal region and is identical to a mutation previously isolated based on its synthetic lethality with the spindle checkpoint mutant *mad1Δ* (Anderson et al., 2007). Thus, *sfi1-R788K* likely directly impairs SPB duplication and/or separation. We also identified several *cdc55* mutations, including *cdc55-G43D* and similar suppression of *spo11Δ spo12Δ* spore viability by *cdc55* mutations has been reported previously (Rabitsch et al., 2003; Kerr et al., 2011). Cdc55 is the B55 regulatory subunit of protein phosphatase 2A, which plays an important role in suppressing release of the phosphatase Cdc14 from the nucleolus in both mitosis and meiosis (Bizzari and Marston, 2011; Queralt et al., 2006). Depletion of Cdc55 during meiosis results in two side-by-side SPBs that fail to separate (Fox et al., 2017), suggesting that the mutations identified here impair SPB separation. A second class of mutations produce a meiosis I spindle, but fail to segregate chromosomes on it. This includes mutations affecting kinetochore function, which likely weaken kinetochore-microtubule interactions. Consistent with this, the Ask1 residue P217 mutated in our screen, lies adjacent to S216, whose Cdk1-dependent phosphorylation is required for proper kinetochore-microtubule attachment strength *in vitro* (Gutierrez et al., 2020). A third class of mutations are likely to have more general effects on cell cycle regulation. Although we did not analyse further here, loss of function mutations in the mitotic/meiotic cyclin Clb4, are known to lead to mis-timed separation of sister chromatids under some circumstances (Lumbroso et al., 2025; Kiburz et al., 2008). Cell cycle kinases, including DDK, Cdc5^Plk1^ and Kns1, also likely disrupt coordination, although their precise mechanisms are less clear.

While the effects of the mutations we identified are subtle in an otherwise wild-type background, they become readily apparent when combined with *spo12Δ*. This indicates that sensitised conditions can expose vulnerabilities in the chromosome segregation machinery that are not evident under normal circumstances, consistent with the broader observation that meiosis can reveal defects that are tolerated during mitosis. For example, components of the Ctf19 kinetochore complex are not essential for mitosis but critical for meiotic kinetochore integrity and chromosome segregation (Borek et al., 2021; Mehta et al., 2014), and loss of the Dam1 complex has similarly disproportionate effects in meiosis in *S. pombe* (Wakiya et al., 2021; Blyth et al., 2018). Likewise, Ase1 plays a particularly important role in *S. pombe* meiosis I (Zheng et al., 2020), as we show here for *S. cerevisiae*. The spindle assembly checkpoint is also less robust during meiosis I (Cairo et al., 2023; Koch et al., 2026), which may further contribute to the consequences of such defects. Furthermore, the mutations predominantly affect meiosis I, while meiosis II and mitosis proceed relatively normally, highlighting the unique sensitivity of this division to kinetochore and spindle perturbations.

### Outlook

The principles revealed here are likely to extend beyond budding yeast. In mammalian oocytes, two successive meiotic divisions occur within a single cytoplasm in the absence of canonical centrosomes, requiring the spindle to be reassembled while preserving the correct chromosome segregation pattern. In this context, the configuration of the chromosomes themselves may provide a key determinant of how they interact with successive spindle cycles. Classic micromanipulation experiments in grasshopper spermatocytes (Paliulis and Nicklas, 2000) and more recent chromosome transplantation studies in mouse oocytes (Ogushi et al., 2021) have demonstrated that chromosome behaviour is largely dictated by their intrinsic state rather than the surrounding cytoplasm. These support a model in which chromosome segregation is coordinated with the spindle cycle through chromosome-intrinsic cues, and suggest that failures in this coupling may contribute to the high rates of segregation errors observed in mammalian meiosis.

## Methods

### Yeast strains and plasmids

All yeast strains used were derivatives of SK1 (*ho::LYS2, ura3, leu2::hisG, trp1::hisG, his3::hisG*) and genotypes are given in Table S2. *pCLB2-CDC20* (Lee and Amon, 2003), *REC8-GFP*, *GFP-TUB1*, *CEN5-tdTomato* (Matos et al., 2008)*, SPC42-tdTomato* (Fox et al., 2017), *spo11Δ::TRP1* (Galander et al., 2019), *hrr25-zo* (Petronczki et al., 2006) were described previously. The plasmids used in the study and details about their creation is in Table S3.

To create the conditional pseudodiploid, a plasmid (AMp1928) was constructed by four-part Gibson assembly using *MATα1–MATα2* amplified from YIPlac211-*MATα* (c4223; (Rabitsch et al., 2003)), *IAA17* amplified from pSM410 (Morawska and Ulrich, 2013), and the vector backbone from YIPlac211-*MATα*. The resulting plasmid was linearized at the *URA3* locus using ApaI and transformed into yeast. The resulting strain was then crossed with a strain generated by transformation with AflII-cut AMp966 (*OsTIR1*) targeted to the *LEU2* locus to generate strain AM29707.

Gene deletions were introduced using PCR-based methods (Longtine et al., 1998). The deletion mutant *spo12 Δ::NAT* and the truncation mutants *ase1Δ473-886 ::KAN, kns1Δ604-737:KANMX,* and *clb4Δ246-460:KANMX* were generated by amplifying a resistance cassette from a plasmid, with homology arms targeting the C-terminus and the gene’s downstream region (for truncations) or only the gene’s flanking regions (for deletions).

*De novo* point mutants to confirm causality were generated by CRISPR–Cas9 genome editing essentially as described in (Borek et al., 2021). Guide RNAs targeting the locus of interest were designed using Benchling and cloned into AMp1278 (pWSO82; (Shaw et al., 2019)) by Golden Gate assembly (Table S2). Yeast were transformed with PCR fragments encoding Cas9, the sgRNA and a homology-directed repair template containing the desired mutation and a modified PAM site to prevent re-cutting. Correctly edited strains were identified by allele-specific PCR and confirmed by Sanger sequencing. Details of construction of individual plasmids are given in Table S3.

### Growth conditions

For sporulation experiments, strains were first grown to saturation in YPDA (1% Bacto-yeast extract, 2% Bacto-peptone, 2% Glucose, 0.3 mM Adenine), diluted to OD_600_ = 0.2 in BYTA medium (1% Bacto-yeast extract, 2% Bacto-tryptone, 1% Potassium acetate, 50mM potassium phthalate), and cultured for 14–16 h at 30 °C with shaking. Cells were then washed twice in water and transferred to sporulation medium (SPO) (0.3% Potassium acetate, pH 7.0) at OD_600_ ≈ 1.8 to induce meiosis. For metaphase I arrest, cells carried *pCLB2-CDC20* (Lee and Amon, 2003).

### Immunoblot analysis

Immunoblotting to confirm Matα-AID degradation was performed as described by (Barton et al., 2022).The primary antibody, anti-AID tag (IAA17) (APC004AM; Cosmo Bio Co. Ltd.; (Nishimura et al., 2009)), was used at 1:500 in 2% milk/PBST overnight, followed by washes in PBST and incubation with HRP-conjugated sheep anti-mouse secondary antibody (GE Healthcare) at 1:5000 in 2% milk/PBST for 1 h at room temperature. Detection and development were performed as described in (Barton et al., 2022).

### Spore viability assay

Yeast spore viability was determined either by standard tetrad dissection or by random spore assay. For tetrad dissection cells were sporulated on agar SPO media at 30°C, suspended in 20 μl 1 mg/ml zymolyase in 2 M sorbitol for 10 min and then diluted with 1 mL water. A minimum of 40 tetrads were dissected into individual spores on YPDA agar using a micromanipulator on a Nikon Eclipse 50i light microscope and viable colonies were scored ∼48 hr later. For random spore assay, sporulated cultures (400 μl of a 2.5 OD_600_ culture or cells from SPO plate) were treated with 1 mg/ml zymolyase in 2 M sorbitol and incubated overnight at 30 °C with rotation to digest asci. Cells were pelleted, washed with water, and resuspended in 10% NP-40 before vigorous vortexing to release spores. The suspension was washed, sonicated (10 cycles of 30s on/30s off, high setting; Bioruptor Plus) to disperse clumps, and spores were counted using a haemocytometer before plating either 200 or 1000 spores on YEPD plates. Colony formation after ∼2 days at 30 °C was used to calculate spore viability.

### EMS mutagenesis and mutant isolation

Random mutagenesis was performed using ethyl methanesulfonate (EMS). Pseudodiploid strain AMy30044 was recovered on YPG plates, transferred to 4% YPD, and inoculated into liquid YPD for overnight growth at 30 °C. The following morning, cells corresponding to 6 OD_600_ units were harvested by centrifugation, washed once with water and resuspended in 1 ml of 0.1 M sodium phosphate buffer (pH 7.4). EMS (15 μl; ∼1.5% v/v final concentration) was added and cells were incubated for 1 h at 30 °C with shaking. Control samples were treated identically but without EMS.

Following mutagenesis, cells were pelleted and resuspended in 200 μl of 5% sodium thiosulfate to inactivate residual EMS, followed by two additional washes with 5% sodium thiosulfate. Cells were then resuspended in 1 ml YEPD and allowed to recover by growth overnight at 30 °C with shaking. Recovered cultures were diluted into BYTA medium to OD_600_ ≈ 0.3 and grown for 14–16 h at 30 °C with shaking. Cells were then washed twice with water and transferred to sporulation medium (SPO) at OD_600_ ≈ 2.5 to induce meiosis. Sporulation cultures were incubated at 30 °C for at least 48 h before performing random spore analysis. Spores were plated on YPDA and incubated at 30 °C for ∼2 days to allow colony formation. Individual colonies were picked and patched onto fresh YPDA plates, and corresponding patches were transferred to SPO plates to assess sporulation phenotypes after 2–3 days. Mutant isolates displaying improved sporulation or spore viability were retained for further analysis and stored as glycerol stocks at −80 °C.

### Mating assay

The cooccurrence of markers unique to individual haploids was used to confirm mating had occurred. MATa and MATα tester strains (Table S1) carried auxotrophic markers absent in SK1 strains (*ho::LYS2, ura3, leu2::hisG, trp1::hisG, his3::hisG*); thus, neither strain grew individually on minimal media (0.67% (w/v) yeast nitrogen base without amino acids, 2% (w/v) glucose, 2% (w/v) agar). SK1 strains were replica-plated onto minimal media plates pre-seeded with MATa or MATα testers and incubated at 30 °C for 2 days. Growth only on MATa tester plates indicated that the SK1 strain was MATα, growth only on MATα tester plates indicated a MATa SK1 strain, and no growth on either tester-seeded minimal plate indicated that the strain was a SK1 diploid or pseudodiploid.

For mating assays that did not use MAT testers (e.g., Fig. S1C, complementation assays, backcrosses, and bulk segregation analysis; see below), mating was assessed by complementation of prototrophic markers in SK1 strains. Strains were mated in liquid YPDA medium containing 500 μM NAA to induce degradation of Matα-AID and permit mating. Mutant and tester strains were mixed at low cell density (final OD₆₀₀ < 0.3) and incubated overnight with gentle rotation. Cultures were then diluted and plated onto selective media to isolate diploids. Double selection was performed on –TRP –LEU plates (0.67% (minimal media supplemented with complete amino acid dropout mix lacking tryptophan and leucine; see Fig. S1C), or by replica plating onto –TRP followed by re-replica plating onto G418 (kanamycin) plates (1% (w/v) yeast extract, 2% (w/v) peptone, 2% (w/v) glucose, 2% (w/v) agar, supplemented with G418 (geneticin) at 250 μg/ml; for complementation assays and backcrosses).

### Complementation assay

Complementation tests were performed by crossing mutant isolates to strains carrying deletions of candidate genes (AMy29865, AMy29871, AMy30030, AMy30965, AMy29951 and AMy31803). To simplify handling of large numbers of mutants, genes were tested in pairs in each round of complementation analysis, typically beginning with *mam1Δ* and *csm1Δ* tester strains. After crosses were setup in liquid media, cultures were diluted and plated onto selective media to isolate diploids. Diploid colonies were patched onto YPD and transferred to sporulation medium (SPO) for 3 days. Complementation was assessed by comparing sporulation phenotypes of diploid crosses to the corresponding parental control strains by light microscopy.

### Backcrossing of mutant isolates

For mutants in which the identity of the gene carrying the causal mutation could not be identified by complementation analysis, isolates were backcrossed to strain AMy29507 in the presence of 500 μM NAA. Multiple independent diploid colonies were patched onto YPD plates before transfer to sporulation medium (SPO) to induce meiosis. Following sporulation, spores were isolated by random spore analysis and at least 10 independent segregants carrying the desired genotype (*MATa spo11Δ::TRP spo12Δ::NAT MATα-aid::URA3 OsTIR1::LEU2*) were isolated and re-tested for the suppressor phenotype by sporulation analysis. Segregants retaining the phenotype were used for subsequent rounds of backcrossing or bulk segregant analysis prior to whole genome sequencing.

### Bulk segregant analysis

Bulk segregant analysis (Gopalakrishnan and Winston, 2019) was used to reduce the contribution of unrelated background mutations introduced during EMS mutagenesis. Following mutagenesis, individual isolates contain numerous mutations. Crossing mutants to a non-mutagenised strain allows these mutations to segregate during meiosis, such that different segregants carry different combinations of background mutations. By selecting multiple segregants that retain the suppressor phenotype and pooling them, the causal mutation shared among them is maintained while unrelated mutations are diluted. This approach provides a rapid alternative to repeated backcrossing.

Each mutant isolate was crossed to strain AMy29507 and diploids were selected by sequential selection for *TRP1* and *KANMX* markers. Diploids were induced to sporulate and spores were isolated by random spore analysis. Approximately 2000 spores were plated and colonies were replica plated onto screening plates (−*URA, −TRP, −LEU, +NAT*) to select the desired haploid progeny. Colonies were patched onto YEPD plates and corresponding patches were transferred to sporulation medium (SPO). Segregants retaining the suppressor phenotype were identified by examining sporulation phenotypes under a light microscope. For each mutant isolate, 20–30 segregants displaying the suppressor phenotype were selected. Individual segregants were first grown separately overnight in 96-well plates. Equal volumes (100 μl) from each culture were then combined and expanded in YEPD in a stepwise manner (15 ml overnight culture followed by expansion to 50 ml) to minimise overrepresentation of individual segregants within the pool. Cells were harvested by centrifugation and stored as pellets for genomic DNA extraction and sequencing.

### Genomic DNA extract for whole-genome sequencing

Genomic DNA was extracted from frozen yeast cell pellets. Cells were resuspended in sorbitol buffer (0.9 M sorbitol, 0.1 M Tris-HCl pH 7.5, 0.1 M EDTA) and treated with zymolyase-100T (10 mg/ml stock; 5 µl per 750 µl suspension) and β-mercaptoethanol (1 µl) for 30 min at 37 °C with rotation to generate spheroplasts. Completion of spheroplasting was verified by lysis in 1% SDS. Spheroplasts were pelleted and resuspended in TE buffer, lysed with 10% SDS, and proteins precipitated by addition of 5 M potassium acetate followed by incubation on ice for 30 min. Lysates were clarified by centrifugation and DNA was precipitated with 2 volumes ethanol. Pellets were resuspended in TE, treated with RNase A (10 mg/ml) for 15 min at 65 °C, and subjected to phenol/chloroform extraction followed by ethanol precipitation with 2 volumes ethanol and 1/10 volume 3 M sodium acetate (pH 5.5). DNA pellets were washed with 70% ethanol, air-dried and resuspended in 100 µl TE. DNA concentrations were determined using Qubit fluorometry. To fragment genomic DNA for sequencing, it was diluted to 10–20 ng/µl in volumes of 100–300 µl and sheared using a Diagenode Bioruptor Plus sonicator (LOW setting, 30 s ON / 30 s OFF for 12 cycles) to generate fragments of ∼400 bp suitable for paired-end sequencing. Fragmentation efficiency was assessed by electrophoresis on a 2% agarose gel.

### Illumina library preparation

Sequencing libraries were prepared using the NEBNext DNA Library Prep Master Mix Set for Illumina with NEXTFLEX adapters. Fragmented DNA was subjected sequentially to end repair, dA-tailing, and adapter ligation following manufacturer recommendations with AMPure XP bead purification between steps. After adapter ligation, libraries were size selected using a two-step AMPure XP bead selection to enrich fragments with an insert size of approximately 400 bp (0.55× bead ratio followed by 0.15×). Libraries were amplified by PCR with an initial denaturation at 98 °C for 30 s, followed by 35 cycles of 98 °C for 10 s and 65 °C for 90 s, and a final extension at 65 °C for 5 min. Amplified libraries were purified using 0.9× AMPure XP beads and eluted in 25 μl TE. Libraries were quantified using the NEBNext Library Quant Kit for Illumina according to the manufacturer’s instructions. Libraries were diluted (10⁻³–10⁻⁵) in dilution buffer and quantified by qPCR using standards ranging from 100 pM to 0.001 pM. Reactions (20 µl) contained 16 µl NEBNext Master Mix and 4 µl template, and were run in triplicate on a Roche LightCycler 480.

### Variant calling pipeline

Genomic DNA from pooled segregants was subjected to Illumina paired-end sequencing. Reads were aligned to the *Saccharomyces cerevisiae* SK1 reference genome (NCBI Accession: PRJEB7245, https://yjx1217.github.io/Yeast_PacBio_2016/welcome) using standard alignment software (Burrows Wheeler Alignment BWA-MEM tool), PCR duplicates removed using a GATK Markduplicates tool, and local realignment was carried out using GATK RealignTargetCreator and GATK IndelRealigner Variants were called using the GATK HaplotypeCaller and written to Variant Call Format (VCF) files Variant annotation and predicted functional effects were determined using SnpEff. Variants were filtered following GATK standard hard-filtering guidelines, including read depth, mapping quality, and strand-bias parameters, and only variants passing all filters were retained for further analysis. Variations common with the unmutagenised parent strain and the strain used for backcrossing was also removed from the analysis. Because sequencing was performed on pooled haploid segregants, genotype calls were less informative than allele frequencies. Candidate causal mutations were therefore identified based on variant allele frequency (VAF), defined as the proportion of reads supporting the alternate allele at a given position (calculated as allelic depth (AD) by total depth (DP) in the VCF file). Mutations linked to the suppressor phenotype were expected to approach fixation within the pool, whereas background mutations segregated randomly and were present at lower allele frequencies.

### Live-cell imaging

Diploid cells were induced to enter meiosis as described above. After 16 h in BYTA medium, cultures were transferred to sporulation (SPO) medium supplemented with 0.2 mM uracil. Cells were incubated at 30 °C with shaking (250 rpm) for 2.5–3 h prior to imaging for both asynchronous and metaphase I arrest cultures. Eight-well glass-bottom Ibidi dishes were coated with 45 μL concanavalin A (ConA) dissolved in 50 mM CaCl₂ and 50 mM MnCl₂ and incubated for 15 min at 30 °C. Wells were washed twice with water before use. Cells (4-5 mL culture) were pelleted at 3000 rpm for 3 min and resuspended in the remaining medium. Approximately 300 μL of cell suspension was added to each well and allowed to adhere for 20 min at 30 °C. Wells were then gently washed twice with water to remove unattached cells and washed once more with conditioned SPO medium from the original culture. After washes 400 μL of conditioned SPO medium was added to each well prior to imaging.

Live-cell imaging was performed at 30 °C using a Zeiss Axio Observer Z1 microscope equipped with a Prior motorised stage and a Hamamatsu Flash4 sCMOS camera. Images were acquired using Zen 2.3 software. For each well, six to seven fields of view were imaged every 15 min for up to 12 h using FITC, tdTomato and brightfield channels. Z-stacks consisting of nine optical sections spaced 0.7 μm apart were collected at each time point. Typical acquisition settings were 200 ms exposure for *CEN5*-tdTomato, 50 ms for GFP–Tub1, 200 ms for Spc42–tdTomato, 50 ms for Rec8-GFP and 10 ms for brightfield imaging, with 2×2 camera binning and 4–10% transmitted light depending on the fluorescent marker.

### Reagent and Data availability

Yeast strains and plasmids generated in this study are available from the lead author, without restriction.

Pipeline for analysis of WGS sequencing data is available on Github with reference: https://github.com/aparna-biologist/Variant_calling_pipeline_AV_SW.git

## Acknowledgements

We gratefully acknowledge the Wellcome Discovery Research Platform for Hidden Cell Biology Light Microscopy and Bioinformatics Cores. We thank Tom Ellis, Kim Nasmyth and the National Bio-Resource Project (NBRP) – Yeast, Japan for plasmids. We are grateful to Weronika Borek for help with initial screen design and to Marston lab members past and present for useful discussion. This work was funded through Wellcome Investigator [220780] and Discovery Awards to AM [319314], a Darwin Trust of Edinburgh studentship to AV, a Wellcome integrated Cell Mechanisms studentship to MT [218470, 316102], core funding for the Wellcome Centre for Cell Biology [203149] and a Wellcome Discovery Research Platform Award [226791].

## Author contributions

Conceptualization – AV and AM; Data curation – AV and SW; Funding Acquisition – AM; Formal Analysis – AV and MT; Investigation – AV and MT; Software – AV and SW; Supervision –AM; Visualization – AM, MT and AV; Writing – original draft – AM; Writing – review and editing – AM, AV and MT.

---

**Table S1 Overview of mutations identified in the screen**

**Table S2 *S. cerevisiae* strains used in this work**

**Table S3 Plasmids generated in this work**

**Figure S1.**
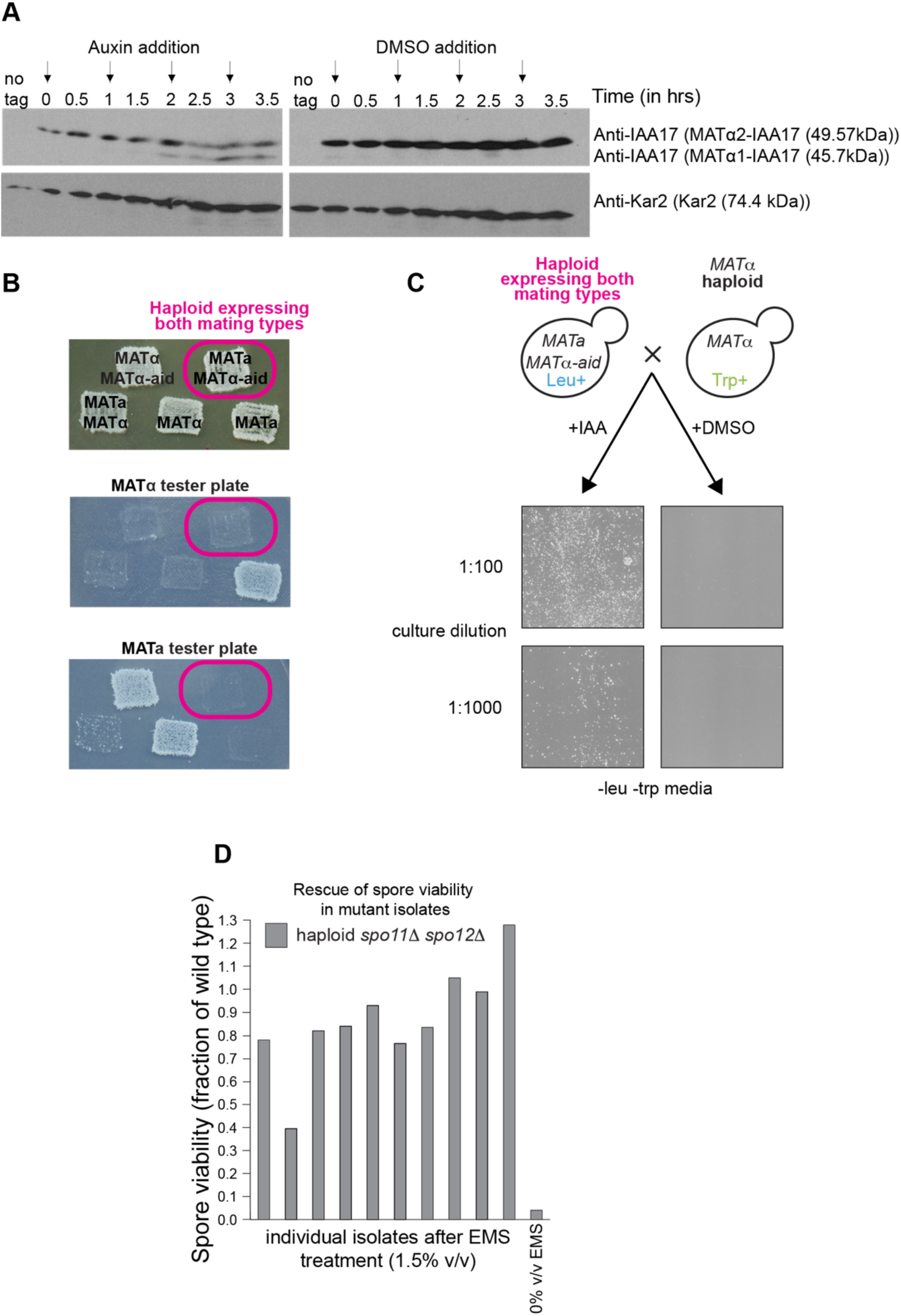
A conditional pseudodiploid. A haploid strain (AM29707) expressing endogenous Mata as well as both Matα factors fused to the auxin-inducible degron (Matα1-aid/Matα2-aid) was constructed. This strain also expresses Tir1, which upon addition of auxin analog (IAA), targets aid to the proteosome. (A) Both Matα1 and Matα2 are degraded in the presence of NAA. Cells were grown in YPDA and samples extracted at the indicated times after addition of 500 μM NAA or DMSO (control) for anti-IAA17 or anti-Kar2 (loading control) western immunoblot analysis. 250 μM NAA, or the same volume of DMSO was re-added every hour (arrows). (B) AM29707 failed to mate with *MATa* or *MATα* tester strains in the absence of NAA. Growth indicates mating due to complementation of auxotrophic markers only upon diploid formation. (C) Treatment of AM29707 with 500μM NAA but not DMSO, allowed mating with a *MATα* strain. Diploid formation is assayed by colony growth on media lacking both leucine and tryptophan, since haploids are individually prototrophic for leucine and tryptophan respectively. (D) Example of spore viability rescue by individual screen isolates after EMS mutagenesis. Viability is presented as a fraction of wild type after random spore assay.

**Figure S2.**
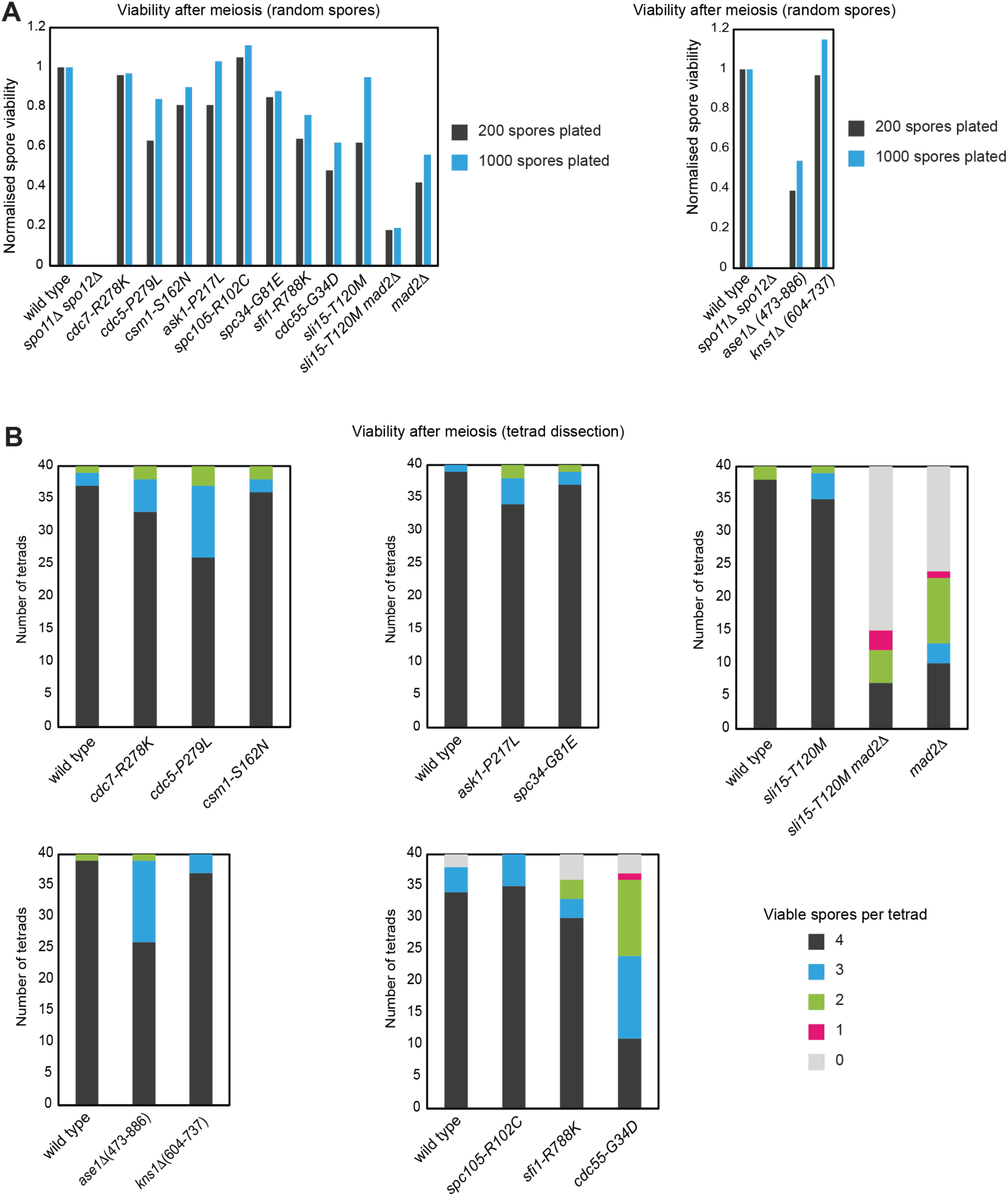
Rescue of spore viability of *spo11Δ spo12Δ* after *de novo* mutant generation. (A) Spore viability is shown normalized to wild type after random spore assay for the indicated *de novo* generated mutants. (B) The fraction of viable spores per tetrad recovered after tetrad dissection is shown for the indicated *de novo* generated mutants.

**Figure S3.**
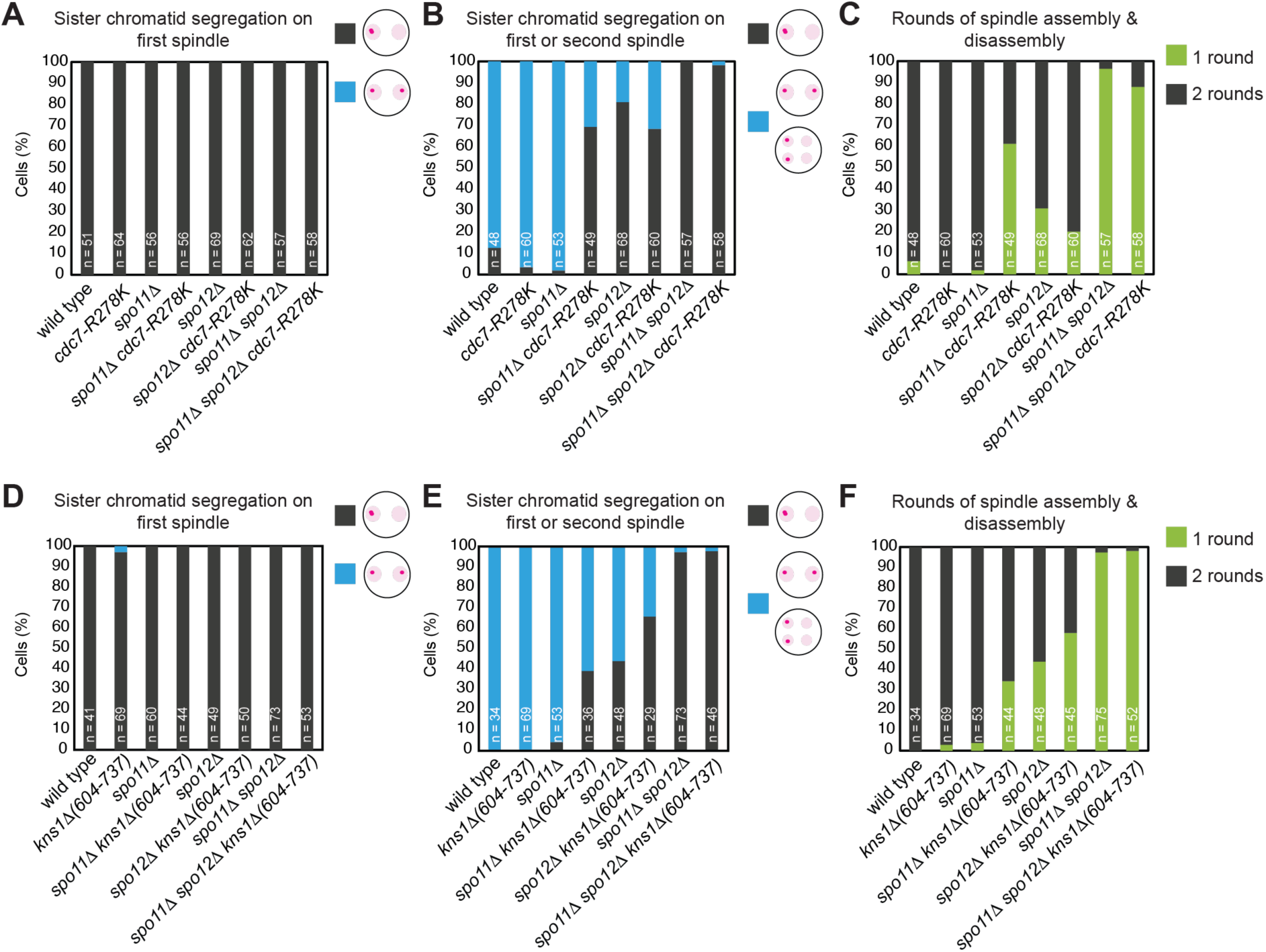
Effect of *cdc7-R278K* and *kns1Δ (604-737)* on sister chromatid segregation. Live imaging as described in Figure 2D-G to examine the effects of *cdc7-R278K* (A-C) or *kns1Δ (604-737)* (D-F). Shown is the percentage of cells where *CEN5-dtTomato* foci separate on the first (A, D) and/or second (B, E) spindle, along with the percentage of cells undergoing one or two rounds of spindle assembly/disassembly (C, F).

**Figure S4.**
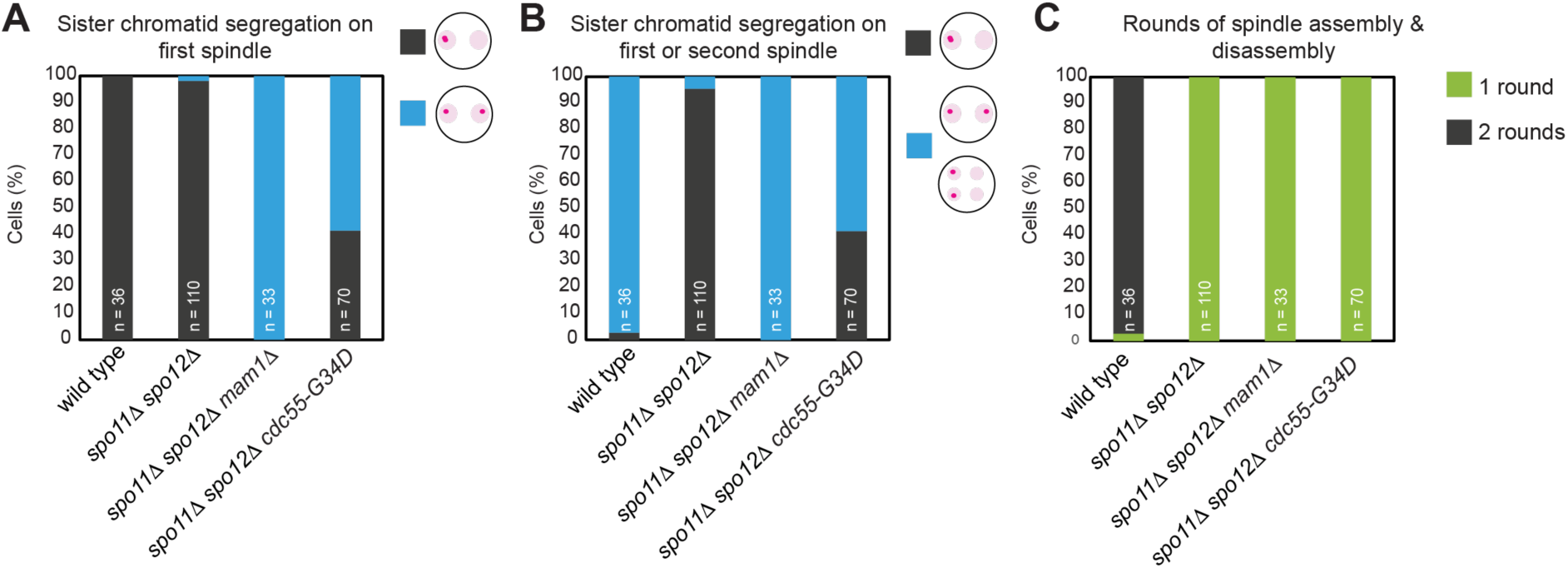
*cdc55-G34D* causes *spo11Δ spo12Δ* cells to undergo only a single round of spindle assembly/disassembly upon which sister chromatids segregate. Live cell imaging as in Figure 2D-G showing the percentage of cells where *CEN5-dtTomato* foci separate on the first (A) and/or second (B) spindle, along with the percentage of cells undergoing one or two rounds of spindle assembly/disassembly (C).

**Figure S5.**
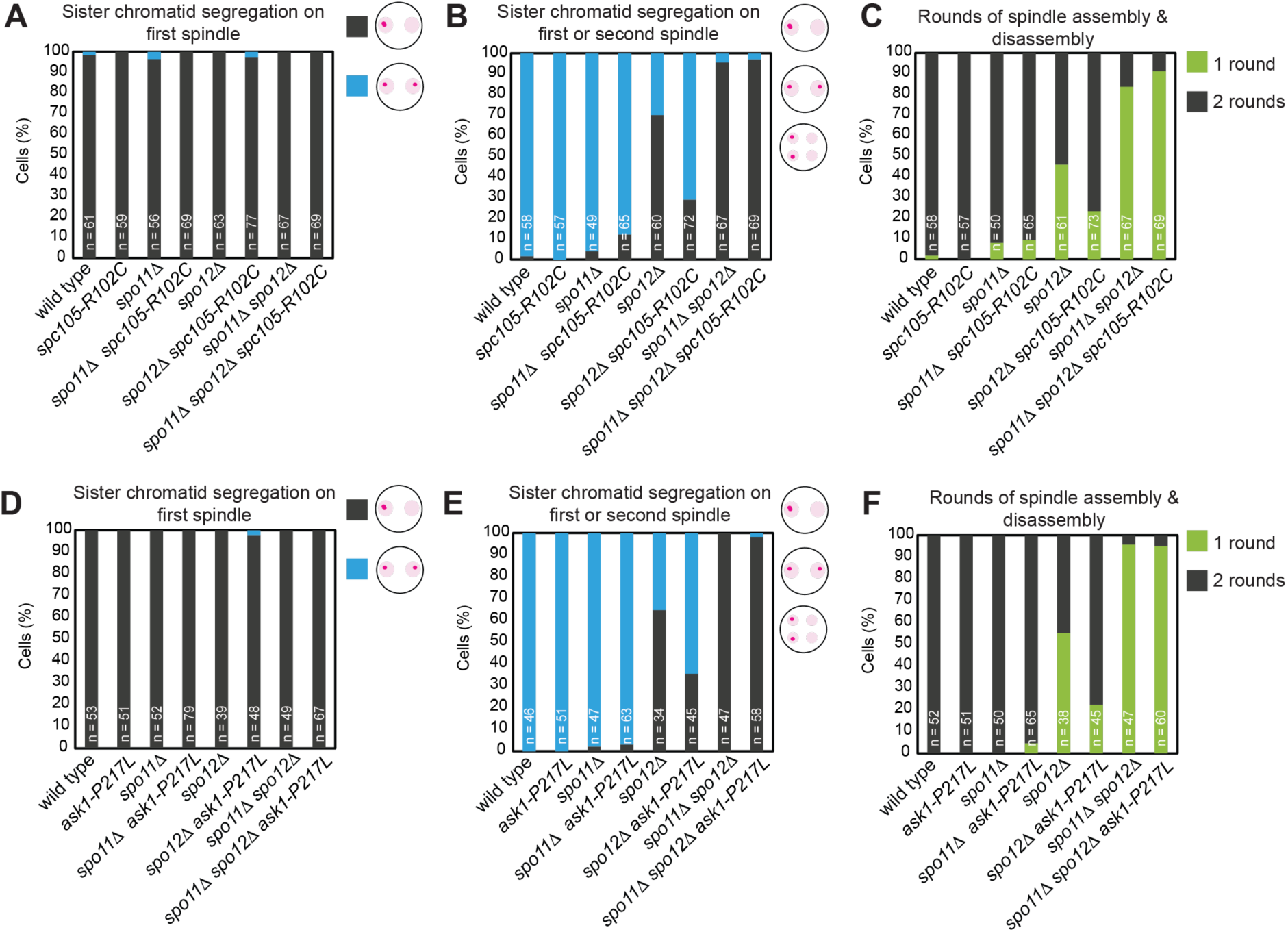
Mutations in kinetochore subunits delay segregation until meiosis II. Live imaging as described in Figure 2D-G to examine the effects of *spc105-R102C* (A-C) or *ask1-P217L* (D-F). Shown is the percentage of cells where *CEN5-dtTomato* foci separate on the first (A, D) and/or second (B, E) spindle, along with the percentage of cells undergoing one or two rounds of spindle assembly/disassembly (C, F).

## References

Anderson, V.E., J. Prudden, S. Prochnik, T.H. Giddings, and K.G. Hardwick. 2007. Novel sfi1 Alleles Uncover Additional Functions for Sfi1p in Bipolar Spindle Assembly and Function. MBoC. 18:2047–2056. doi:10.1091/mbc.e06-10-0918.

Avena, J.S., S. Burns, Z. Yu, C.C. Ebmeier, W.M. Old, S.L. Jaspersen, and M. Winey. 2014. Licensing of Yeast Centrosome Duplication Requires Phosphoregulation of Sfi1. PLOS Genetics. 10:e1004666. doi:10.1371/journal.pgen.1004666.

Barton, R.E., L.F. Massari, D. Robertson, and A.L. Marston. 2022. Eco1-dependent cohesin acetylation anchors chromatin loops and cohesion to define functional meiotic chromosome domains. Elife. 11:e74447. doi:10.7554/eLife.74447.

Bizzari, F., and A.L. Marston. 2011. Cdc55 coordinates spindle assembly and chromosome disjunction during meiosis. Journal of Cell Biology. 193:1213–1228. doi:10.1083/jcb.201103076.

Blyth, J., V. Makrantoni, R.E.. Barton, C. Spanos, J. Rappsilber, and A.L.A.L. Marston. 2018. Genes important for schizosaccharomyces pombe meiosis identified through a functional genomics screen. Genetics. 208:589–603. doi:10.1534/genetics.117.300527.

Borek, W.E., N. Vincenten, E. Duro, V. Makrantoni, C. Spanos, K.K. Sarangapani, F. de Lima Alves, D.A. Kelly, C.L. Asbury, J. Rappsilber, and A.L. Marston. 2021. The Proteomic Landscape of Centromeric Chromatin Reveals an Essential Role for the Ctf19CCAN Complex in Meiotic Kinetochore Assembly. Curr Biol. 31:283–296.e7. doi:10.1016/j.cub.2020.10.025.

Briza, P., E. Bogengruber, A. Thür, M. Rützler, M. Münsterkötter, I.W. Dawes, and M. Breitenbach. 2002. Systematic analysis of sporulation phenotypes in 624 non-lethal homozygous deletion strains of Saccharomyces cerevisiae. Yeast. 19:403–422. doi:10.1002/yea.843.

Buonomo, S.B.C., K.P. Rabitsch, J. Fuchs, S. Gruber, M. Sullivan, F. Uhlmann, M. Petronczki, A. Tóth, and K. Nasmyth. 2003. Division of the nucleolus and its release of CDC14 during anaphase of meiosis I depends on separase, SPO12, and SLK19. Developmental cell. 4:727–739.

Cairo, G., C. Greiwe, G.I. Jung, C. Blengini, K. Schindler, and S. Lacefield. 2023. Distinct Aurora B pools at the inner centromere and kinetochore have different contributions to meiotic and mitotic chromosome segregation. Mol Biol Cell. 34:ar43. doi:10.1091/mbc.E23-01-0014.

Chelysheva, L., S. Diallo, D. Vezon, G. Gendrot, N. Vrielynck, K. Belcram, N. Rocques, A. Marquez-Lema, A.M. Bhatt, C. Horlow, R. Mercier, C. Mezard, and M. Grelon. 2005. AtREC8 and AtSCC3 are essential to the monopolar orientation of the kinetochores during meiosis. J Cell Sci. 118:4621–4632.

Chen, Y.-C., and M. Weinreich. 2010. Dbf4 regulates the Cdc5 Polo-like kinase through a distinct non-canonical binding interaction. J Biol Chem. 285:41244–41254.

Clyne, R.K., V.L. Katis, L. Jessop, K.R. Benjamin, I. Herskowitz, M. Lichten, and K. Nasmyth. 2003. Polo-like kinase Cdc5 promotes chiasmata formation and cosegregation of sister centromeres at meiosis I. Nat Cell Biol. 5:480–485.

Corbett, K.D., and S.C. Harrison. 2016. Molecular Architecture of the Yeast Monopolin Complex. Cell Rep. 17:929. doi:10.1016/j.celrep.2016.09.070.

Corbett, K.D., C.K. Yip, L.-S. Ee, T. Walz, A. Amon, and S.C. Harrison. 2010. The monopolin complex crosslinks kinetochore components to regulate chromosome-microtubule attachments. Cell. 142:556–567. doi:10.1016/j.cell.2010.07.017.

Duro, E., and A.L. Marston. 2015. From equator to pole: splitting chromosomes in mitosis and meiosis. Genes Dev. 29:109–122. doi:10.1101/gad.255554.114.

Elserafy, M., M. Šarić, A. Neuner, T. Lin, W. Zhang, C. Seybold, L. Sivashanmugam, and E. Schiebel. 2014. Molecular Mechanisms that Restrict Yeast Centrosome Duplication to One Event per Cell Cycle. Current Biology. 24:1456–1466. doi:10.1016/j.cub.2014.05.032.

Enyenihi, A.H., and W.S. Saunders. 2003. Large-scale functional genomic analysis of sporulation and meiosis in Saccharomyces cerevisiae. Genetics. 163:47–54.

Esposito, M.S., R.E. Esposito, M. Arnaud, and H.O. Halvorson. 1970. Conditional mutants of meiosis in yeast. J Bacteriol. 104:202–210. doi:10.1128/jb.104.1.202-210.1970.

Fox, C., J. Zou, J. Rappsilber, and A.L.A.L. Marston. 2017. Cdc14 phosphatase directs centrosome re-duplication at the meiosis I to meiosis II transition in budding yeast. Wellcome open research. 2:2. doi:10.12688/wellcomeopenres.10507.1.

Galander, S., R.E. Barton, W.E. Borek, C. Spanos, D.A. Kelly, D. Robertson, J. Rappsilber, and A.L. Marston. 2019. Reductional Meiosis I Chromosome Segregation Is Established by Coordination of Key Meiotic Kinases. Dev Cell. 49:526–541.e5. doi:10.1016/j.devcel.2019.04.003.

Gopalakrishnan, R., and F. Winston. 2019. Whole genome sequencing of yeast cells. Curr Protoc Mol Biol. 128:e103. doi:10.1002/cpmb.103.

Gutierrez, A., J. ook Kim, N.T. Umbreit, C.L. Asbury, T.N. Davis, M.P. Miller, and S. Biggins. 2020. Cdk1 Phosphorylation of the Dam1 Complex Strengthens Kinetochore-Microtubule Attachments. Current Biology. 30:4491–4499.e5. doi:10.1016/j.cub.2020.08.054.

Katis, V.L., J. Matos, S. Mori, K. Shirahige, W. Zachariae, and K. Nasmyth. 2004. Spo13 facilitates monopolin recruitment to kinetochores and regulates maintenance of centromeric cohesion during yeast meiosis. Curr Biol. 14:2183–2196. doi:10.1016/j.cub.2004.12.020.

Keeney, S., C.N. Giroux, and N. Kleckner. 1997. Meiosis-specific DNA double-strand breaks are catalyzed by Spo11, a member of a widely conserved protein family. Cell. 88:375–384.

Kerr, G.W., S. Sarkar, K.L. Tibbles, M. Petronczki, J.B.A. Millar, and P. Arumugam. 2011. Meiotic nuclear divisions in budding yeast require PP2A(Cdc55)-mediated antagonism of Net1 phosphorylation by Cdk. The Journal of cell biology. 193:1157–1166.

Kiburz, B.M.M., A. Amon, and A.L.L. Marston. 2008. Shugoshin Promotes Sister Kinetochore Biorientation in Saccharomyces cerevisiae. Molecular biology of the cell. 19:1199–1209. doi:10.1091/mbc.E07-06-0584.

Kim, J., K.-I. Ishiguro, A. Nambu, B. Akiyoshi, S. Yokobayashi, A. Kagami, T. Ishiguro, A.M. Pendás, N. Takeda, Y. Sakakibara, T.S. Kitajima, Y. Tanno, T. Sakuno, and Y. Watanabe. 2015. Meikin is a conserved regulator of meiosis-I-specific kinetochore function. Nature. 517:466–471.

Klapholz, S., and R.E. Esposito. 1980. Isolation of SPO12-1 and SPO13-1 from a natural variant of yeast that undergoes a single meiotic division. Genetics. 96:567–588.

Koch, L.B., T. Ghosh, C. Spanos, and A.L. Marston. 2026. Specialisation of meiotic kinetochores revealed through a synthetic spindle assembly checkpoint strategy. Elife. 15:RP110117. doi:10.7554/eLife.110117.

Lee, B.H., and A. Amon. 2003. Role of Polo-like kinase CDC5 in programming meiosis I chromosome segregation. Science (New York, N.Y.). 300:482–486.

Lee, B.H., B.M. Kiburz, and A. Amon. 2004. Spo13 maintains centromeric cohesion and kinetochore coorientation during meiosis I. Curr Biol. 14:2168–2182. doi:10.1016/j.cub.2004.12.033.

Longtine, M.S., A. 3rd McKenzie, D.J. Demarini, N.G. Shah, A. Wach, A. Brachat, P. Philippsen, and J.R. Pringle. 1998. Additional modules for versatile and economical PCR-based gene deletion and modification in Saccharomyces cerevisiae. Yeast. 14:953–961.

Lumbroso, G., G. Cairo, S. Lacefield, and A.W. Murray. 2025. The B-type cyclin Clb4 prevents meiosis I sister centromere separation in budding yeast. G3 (Bethesda). 15:jkaf121. doi:10.1093/g3journal/jkaf121.

Malone, R.E., S. Bullard, M. Hermiston, R. Rieger, M. Cool, and A. Galbraith. 1991. Isolation of mutants defective in early steps of meiotic recombination in the yeast Saccharomyces cerevisiae. Genetics. 128:79–88. doi:10.1093/genetics/128.1.79.

Marston, A.L., B.H. Lee, and A. Amon. 2003. The Cdc14 phosphatase and the FEAR network control meiotic spindle disassembly and chromosome segregation. Developmental cell. 4:711–726. doi:10.1016/S1534-5807(03)00130-8.

Marston, A.L.L., W.-H.H. Tham, H. Shah, and A. Amon. 2004. A genome-wide screen identifies genes required for centromeric cohesion. Science (New York, N.Y.). 303:1367–1370. doi:10.1126/science.1094220.

Matos, J., J.J. Lipp, A. Bogdanova, S. Guillot, E. Okaz, M. Junqueira, A. Shevchenko, and W. Zachariae. 2008. Dbf4-dependent CDC7 kinase links DNA replication to the segregation of homologous chromosomes in meiosis I. Cell. 135:662–678.

Mehta, G.D., M. Agarwal, and S.K. Ghosh. 2014. Functional characterization of kinetochore protein, Ctf19 in meiosis I: An implication of differential impact of Ctf19 on the assembly of mitotic and meiotic kinetochores in Saccharomyces cerevisiae. Molecular Microbiology. 91:1179–1199. doi:10.1111/mmi.12527.

Mengoli, V., K. Jonak, O. Lyzak, M. Lamb, L.M. Lister, C. Lodge, J. Rojas, I. Zagoriy, M. Herbert, and W. Zachariae. 2021. Deprotection of centromeric cohesin at meiosis II requires APC/C activity but not kinetochore tension. The EMBO journal. 40. doi:10.15252/EMBJ.2020106812.

Morawska, M., and H.D. Ulrich. 2013. An expanded tool kit for the auxin-inducible degron system in budding yeast. Yeast. 30:341–351. doi:10.1002/yea.2967.

Mukherjee, A., C. Spanos, and A.L. Marston. 2024. Distinct roles of spindle checkpoint proteins in meiosis. Curr Biol. 34:3820–3829.e5. doi:10.1016/j.cub.2024.07.025.

Nairz, K., and F. Klein. 1997. mre11S--a yeast mutation that blocks double-strand-break processing and permits nonhomologous synapsis in meiosis. Genes & development. 11:2272–2290.

Nilsson, J. 2018. Protein phosphatases in the regulation of mitosis. J Cell Biol. 218:395–409. doi:10.1083/jcb.201809138.

Nishimura, K., T. Fukagawa, H. Takisawa, T. Kakimoto, and M. Kanemaki. 2009. An auxin-based degron system for the rapid depletion of proteins in nonplant cells. Nat Methods. 6:917–922.

Ogushi, S., A. Rattani, J. Godwin, J. Metson, L. Schermelleh, and K. Nasmyth. 2021. Loss of sister kinetochore co-orientation and peri-centromeric cohesin protection after meiosis I depends on cleavage of centromeric REC8. Dev Cell. 56:3100–3114.e4. doi:10.1016/j.devcel.2021.10.017.

Padmanabha, R., S. Gehrung, and M. Snyder. 1991. The KNS1 gene of Saccharomyces cerevisiae encodes a nonessential protein kinase homologue that is distantly related to members of the CDC28/cdc2 gene family. Mol Gen Genet. 229:1–9. doi:10.1007/BF00264206.

Paliulis, L.V., and R.B. Nicklas. 2000. The reduction of chromosome number in meiosis is determined by properties built into the chromosomes. The Journal of cell biology. 150:1223–1231. doi:10.1083/JCB.150.6.1223.

Parra, M.T., A. Viera, R. Gomez, J. Page, R. Benavente, J.L. Santos, J.S. Rufas, and J.A. Suja. 2004. Involvement of the cohesin Rad21 and SCP3 in monopolar attachment of sister kinetochores during mouse meiosis I. J Cell Sci. 117:1221–1234.

Petronczki, M., J. Matos, S. Mori, J. Gregan, A. Bogdanova, M. Schwickart, K. Mechtler, K. Shirahige, W. Zachariae, and K. Nasmyth. 2006. Monopolar attachment of sister kinetochores at meiosis I requires casein kinase 1. Cell. 126:1049–1064.

Plowman, R., N. Singh, E.C. Tromer, A. Payan, E. Duro, C. Spanos, J. Rappsilber, B. Snel, G.J.P.L. Kops, K.D. Corbett, and A.L. Marston. 2019. The molecular basis of monopolin recruitment to the kinetochore. Chromosoma. 128:331–354. doi:10.1007/s00412-019-00700-0.

Queralt, E., C. Lehane, B. Novak, and F. Uhlmann. 2006. Downregulation of PP2A(Cdc55) phosphatase by separase initiates mitotic exit in budding yeast. Cell. 125:719–732.

Rabitsch, K.P., M. Petronczki, J.-P. Javerzat, S. Genier, B. Chwalla, A. Schleiffer, T.U. Tanaka, and K. Nasmyth. 2003. Kinetochore recruitment of two nucleolar proteins is required for homolog segregation in meiosis I. Developmental cell. 4:535–548.

Rüthnick, D., and E. Schiebel. 2016. Duplication of the Yeast Spindle Pole Body Once per Cell Cycle. Mol Cell Biol. 36:1324–1331.

Sakuno, T., K. Tada, and Y. Watanabe. 2009. Kinetochore geometry defined by cohesion within the centromere. Nature. 458:852–858.

Sarangapani, K.K., E. Duro, Y. Deng, F. De Lima Alves, Q. Ye, K.N. Opoku, S. Ceto, J. Rappsilber, K.D. Corbett, S. Biggins, A.L. Marston, and C.L. Asbury. 2014. Sister kinetochores are mechanically fused during meiosis i in yeast. Science. 346:248–251. doi:10.1126/science.1256729.

Sarkar, S., R.T. Shenoy, J.Z. Dalgaard, L. Newnham, E. Hoffmann, J.B.A. Millar, and P. Arumugam. 2013. Monopolin subunit Csm1 associates with MIND complex to establish monopolar attachment of sister kinetochores at meiosis I. PLoS genetics. 9:e1003610.

Severson, A.F., L. Ling, V. Van Zuylen, and B.J. Meyer. 2009. The axial element protein HTP-3 promotes cohesin loading and meiotic axis assembly in C. elegans to implement the meiotic program of chromosome segregation. Genes and Development. 23:1763–1778. doi:10.1101/gad.1808809.

Shaw, W.M., H. Yamauchi, J. Mead, G.O.F. Gowers, D.J. Bell, D. Öling, N. Larsson, M. Wigglesworth, G. Ladds, and T. Ellis. 2019. Engineering a Model Cell for Rational Tuning of GPCR Signaling. Cell. 177:782–796.e27. doi:10.1016/j.cell.2019.02.023.

Shonn, M.A., R. McCarroll, and A.W. Murray. 2000. Requirement of the spindle checkpoint for proper chromosome segregation in budding yeast meiosis. Science (New York, N.Y.). 289:300–303.

Singh, D.-K., A. Mahlandt, S. Jolivet, S. Durand, B. Walkemeier, C. Taochy, V. Solier, M. Derkacheva, L. Cromer, and R. Mercier. 2025. Monopolar orientation of kinetochores at meiosis is enforced by COHESINS and their regulators, CENP-C, and the deSUMOylase SPF2. 2025.08.19.671082. doi:10.1101/2025.08.19.671082.

Toth, A., K.P. Rabitsch, M. Galova, A. Schleiffer, S.B. Buonomo, and K. Nasmyth. 2000. Functional genomics identifies monopolin: a kinetochore protein required for segregation of homologs during meiosis i. Cell. 103:1155–1168.

Tsuboi, M. 1983. The isolation and genetic analysis of sporulation-deficient mutants in Saccharomyces cerevisiae. Mol Gen Genet. 191:17–21. doi:10.1007/BF00330883.

Wakiya, M., E. Nishi, S. Kawai, K. Yamada, K. Katsumata, A. Hirayasu, Y. Itabashi, and A. Yamamoto. 2021. Chiasmata and the kinetochore component Dam1 are crucial for elimination of erroneous chromosome attachments and centromere oscillation at meiosis I. Open Biol. 11:200308. doi:10.1098/rsob.200308.

Ye, Q., S.N. Ur, T.Y. Su, and K.D. Corbett. 2016. Structure of the Saccharomyces cerevisiae Hrr25:Mam1 monopolin subcomplex reveals a novel kinase regulator. Embo J. 35:2139–2151.

Yokobayashi, S., and Y. Watanabe. 2005. The kinetochore protein Moa1 enables cohesion-mediated monopolar attachment at meiosis I. Cell. 123:803–817. doi:10.1016/j.cell.2005.09.013.

Zheng, F., F. Dong, S. Yu, T. Li, Y. Jian, L. Nie, and C. Fu. 2020. Klp2 and Ase1 synergize to maintain meiotic spindle stability during metaphase I. Journal of Biological Chemistry. 295:13287–13298. doi:10.1074/jbc.RA120.012905.

